# Photoaffinity enabled transcriptome-wide identification of splice modulating small molecule-RNA binding events in native cells

**DOI:** 10.1101/2024.11.14.623654

**Authors:** Raven Shah, Wanlin Yan, Joyce Rigal, Steve Mullin, Lin Fan, Lynn McGregor, Andrew Krueger, Nicole Renaud, Andrea Byrnes, Jason R. Thomas

## Abstract

Splice modulating small molecules have been developed to promote the U1 snRNP to engage with pre-mRNAs with strong and altered sequence preference. Transcriptomic profiling of bulk RNA from compound treated cells enables detection of RNAs impacted; however, it is difficult to delineate whether transcriptional changes are a consequence of direct compound treatment or *trans*-acting effects. To identify RNA targets that bind directly with splice modulating compounds, we deployed a photoaffinity labeling (PAL)-based Chem-CLIP approach. Through this workflow, we identify the telomerase lncRNA (TERC) as a previously unknown target of this class of clinically relevant small molecules. Using cellular Δ SHAPE-MaP, we orthogonally validate and further define the compound binding site as likely to be the conserved CR4/5 domain. Additionally, a thorough analysis of the PAL-based Chem-CLIP data reveals that considering competed RNAs, irrespective of magnitude of enrichment, adds a rich dimension of hit calling.

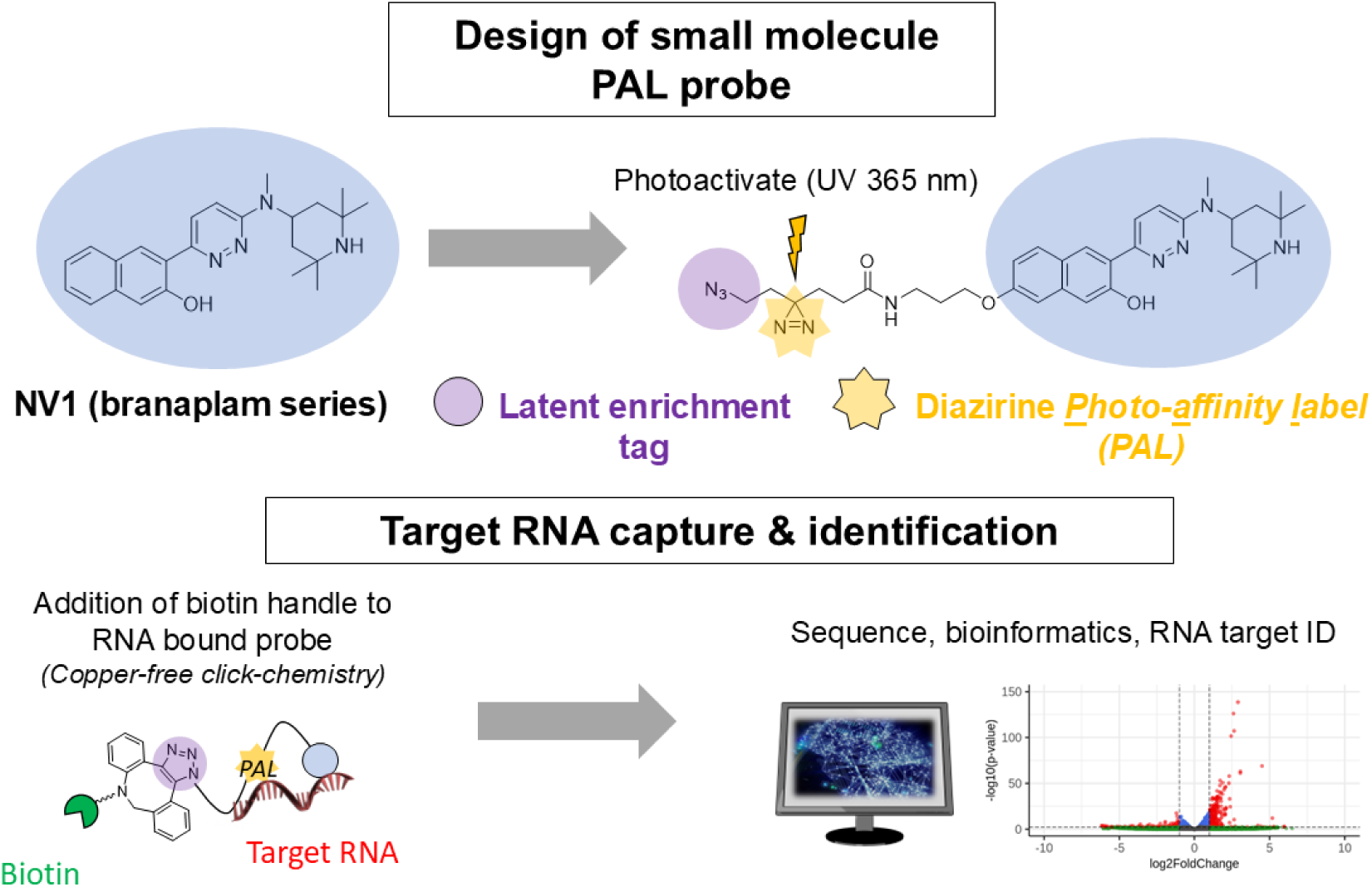

## Introduction

Splicing modulation via RNA-targeting small molecules is an established effective therapeutic modality in the treatment of neurodegenerative diseases such as spinal muscular atrophy (SMA) and Huntington’s disease (HD) ^1–7^. These splicing-modulating compounds interact directly with pre-mRNA/U1snRNP complexes to induce therapeutically favorable changes in splicing to alter disease progression. In the case of SMA, exclusion of exon 7 in the *SMN2* gene prevents canonical RNA transcript expression, leading to an aberrant isoform that is quickly degraded through nonsense mediated decay pathways ^1, 2, 4^. Treatment with branaplam and risdiplam (Evrysdi®) can promote exon 7 inclusion by enhancing the engagement of U1snRNP with the nonconical GGA 3’ exon ending motif^1, 2, 4^.

Branaplam and structurally related analogs show therapeutic efficacy in the treatment of HD, which is caused by CAG trinucleotide expansion repeats in exon 1 of the *HTT* gene. CAG repeat expansion leads to mutant HTT protein aggregation and downstream progressive neurodegeneration ^2, 7^. Splicing modulation can deplete mutant *HTT* mRNA via inclusion of a “poison” cryptic exon that leads to frameshift-induced rapid transcript degradation ^2, 7^. Other recent expression profiling studies with branaplam and risdiplam treatment have identified additional compound-induced cryptic exons in disease-relevant mRNAs, where inclusion of these exons led to significant reduction in transcript levels. These findings highlight the potential to use RNA-interacting small molecules to regulate target RNA steady-state levels to treat a broad spectrum of genetic disorders ^8–10^.

Branaplam and risdiplam function as “molecular glues” bringing together the splicing machinery to nGA 3’ exonic ends in an U1 snRNP-dependent manner ^1, 2^. Within the context of SMN2, proposed mechanisms suggest that branaplam and risdiplam act at an unpaired adenosine bulge at the –1 position (−1A) of the pre-mRNA splice site ^3, 11^. Recent single-molecule imaging studies not only validate the –1A bulge hypothesis but also suggest that branaplam alters U1 snRNP splice site recognition. This occurs through a sequential binding mechanism where compounds bind to the U1 snRNP/U1-C complex after the initial engagement and stabilization of the –1A bulged residue at the splice junction^12^. Time-dependent transcriptomic profiling of branaplam and risdiplam suggest these compounds have diverse target profiles that fluctuate across treatment duration. Though, it remains to be understood whether the extent to which these transcriptional fluctuations are a consequence of direct compound engagement or of splicing events that have already occurred ^2, 11, 13^.

Capturing direct engagement between small molecule and their RNA targets can provide improved insight into compound selectivity, RNA/RNP ligandability, and functional nodes of compound regulation. The applications of electrophilic or PAL moieties, widely used in the areas of covalent drug discovery and chemoproteomics ^14–17^, have been adapted to enable covalent enrichment of small molecule bound RNAs coupled with next-generation sequencing (NGS) ^18–22^. To identify the RNA interactors for the branaplam series of splicing modulators, we adopted a PAL-based Chem-CLIP workflow, as this strategy has been successfully applied to capture small molecule-RNA engagement in a PreQ_1_ riboswitch system ^20^ and RNA interactors of a dihydropyrimidine 1 photoaffinity probe ^19^. Our aim was to establish a workflow for profiling transcriptome-wide compound-RNA binding events in native cellular environments using mechanistically well-understood tool compounds.

## Results and Discussion

### Bioactivity characterization of diazirine-based PAL chemical probe

A photoaffinity probe based on **NV1** was designed to incorporate a photocrosslinkable diazirine moiety and a terminal azide to enable crosslinking and subsequent enrichment (**Fig 1(a)**). Compound bioactivity and cytotoxicity were assessed using a stably integrated *SMN2* luciferase mini-gene reporter SH-SY5Y cell line. The reporter construct design and readout were as previously reported^1^. In this cell model, corrective splicing induction of exon7 within the SMN2 cassette is monitored *via* luciferase activity. The **NV1**-based PAL probe, **NV1-PAL**, retains comparable splicing activity as **NV1** (**Fig 1(b)**). Also, no statistically significant change in cell viability was observed with up to 10 µM **NV1-PAL** treatment for 24 hr (**Fig 1(c)**). Splicing activity at the *SMN2* genetic locus was comparable for **NV1** and **NV1-PAL** at 1 µM treatment at the 3 hr mark (**Fig.1(d)**). Additionally, we confirmed that **NV1-PAL** retained the ability to crosslink with target RNA in cells (**Fig 1(e)**). Confident that **NV1-PAL** maintained splicing modulating activity and is capable of photocrosslinking to RNA targets, we next set out to apply a streamlined workflow for unbiased profiling of small molecule RNA interactions in cells.

**Figure 1:**
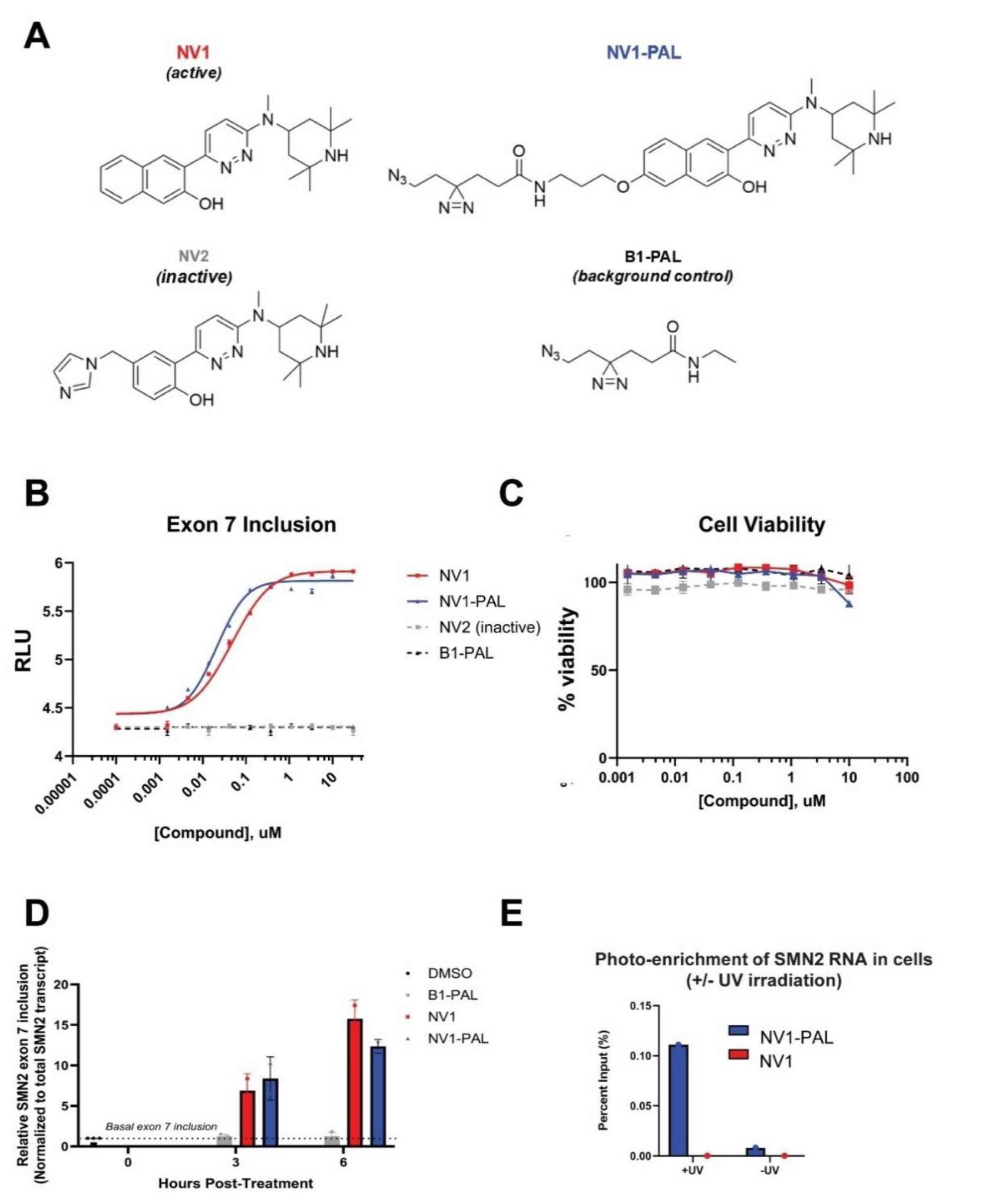
Structures and functional characterization of compounds used in this study. (A) Structures of compounds used in this study: **NV1** (active in SMN2 mini-gene assay), **NV2** (inactive in SMN2 mini-gene assay), **NV1-PAL,** and **B1-PAL**. (B) Dose-response curves of SMN2 luciferase mini-gene reporter stably integrated into SH-SY5Y cells. (C) Dose-response CTG viability assay. Both SMN2 mini-gene and CTG viability assays were run in parallel with 24 hr readout. (D) Kinetics of compound-induced splicing of SMN2 reporter gene as determined by RT-qPCR. Cells were treated with 1 µM of indicated compound. Relative SMN2 exon 7 inclusion was normalized to total SMN2 reporter transcript levels. (E) Photocrosslinking of SMN2 reporter gene. Enrichment of **NV1-PAL** *vs.* **NV1** was assessed without or with UV irradiation using a RT-qPCR read-out following streptavidin enrichment. Degree of enrichment was normalized to total input prior to streptavidin enrichment.

### Applying PAL-based Chem-CLIP in a native cellular environment to identify putative RNA targets

To enable transcriptome-wide profiling of endogenous RNA-small molecule interactions, we applied a PAL-based Chem-CLIP workflow as represented schematically in **Fig 2**. Consistent with other studies ^20, 21^, two criteria were selected for identifying RNAs likely to be direct interactors of **NV1**: (1) a target RNA should be enriched by **NV1-PAL** and (2) the enrichment of that RNA should be diminished by pre-treatment with excess **NV1**; i.e., competition. We multiplexed PAL-based Chem-CLIP samples with identically and contemporaneously treated samples for bulk transcriptomic profiling. We anticipated that this additional layer of multiplexing would enable correlations of enrichment for **NV1-PAL**-labeled RNAs with changes in expression or splicing. Also, this additional layer of multiplexing would enable direct assessment of PAL enriched transcripts with their abundance prior to enrichment across all treatment conditions.

**Figure 2:**
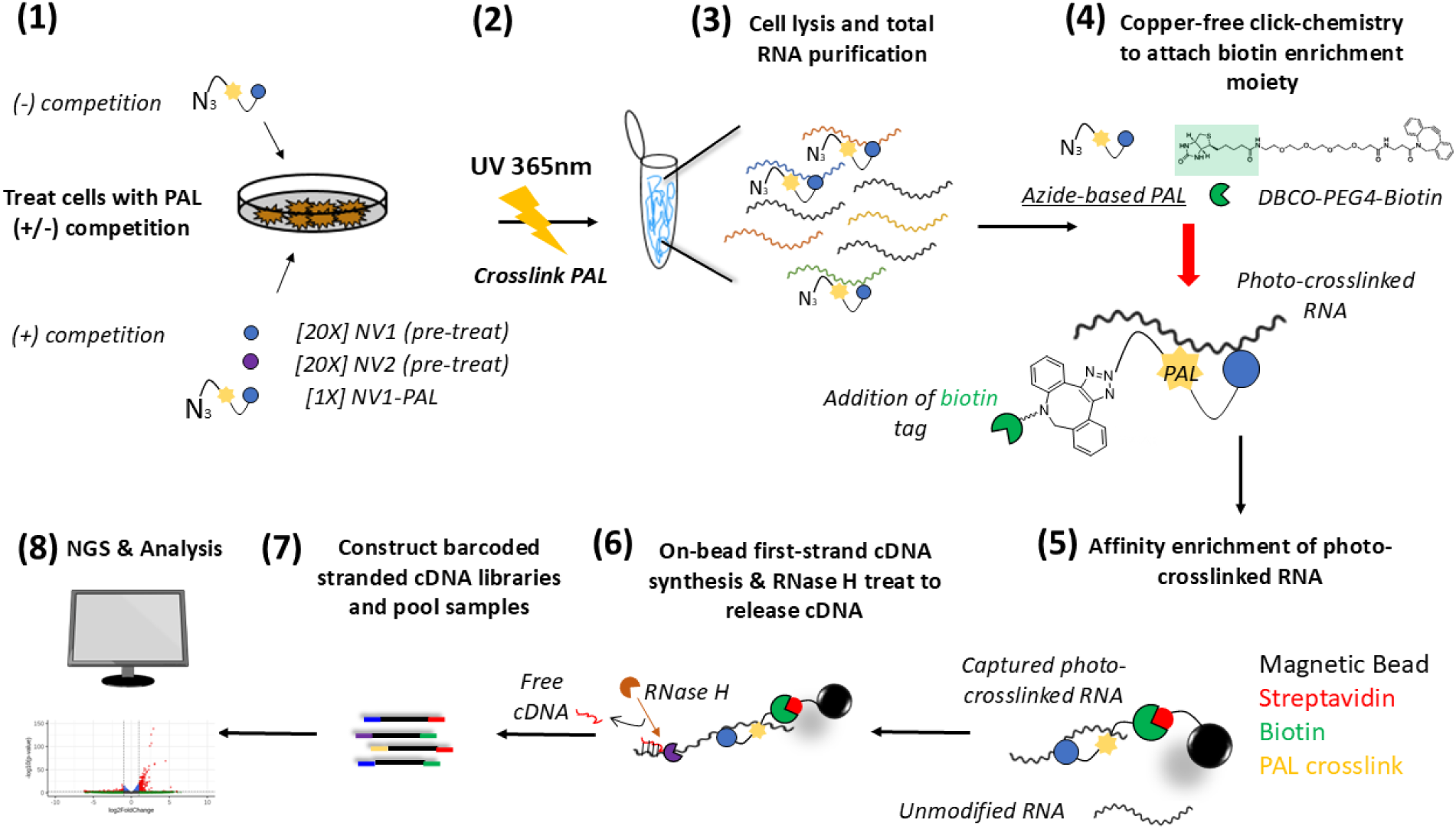
PAL-based chem-CLIP: treatment conditions and workflow schematic. (1) Wildtype SH-SY5Y cells are treated with 1 µM PAL probe (**NV1-PAL** or **B1-PAL**). For competition samples, cells were pre-treated with 20 µM **NV1** or 20 µM **NV2** for 1 hr prior to **NV1-PAL** probe addition. (2) Photocrosslinking is initiated by exposure to UV light (365 nm) for 20 min. (3) Cell lysis and total RNA purification. (4) Copper-free click chemistry to attach biotin for RNA enrichment and clean-up total RNA. (5) Enrichment of photocrosslinked RNA using magnetic streptavidin beads. (6) On-bead first-strand cDNA synthesis and RNase H treat to releases cDNA. (7) Construct barcoded stranded cDNA libraries and pooled. (8) NGS sequencing.

Native, wildtype SH-SY5Y cells were initially treated with DMSO, 20 µM **NV1,** or 20 µM **NV2** for 1 hr. This pre-treatment step allows the compounds to occupy binding sites on target RNAs and thus, prevent subsequent PAL enrichment. After pre-treatment, cells are then treated with 1 µM **NV1-PAL** for 3 hrs. To distinguish enrichment of RNAs driven by **NV1** scaffold from enrichment driven by the PAL moiety, cells were also treated with 1 µM **B1-PAL** probe. Upon UV irradiation, the diazirine is converted into a carbene intermediate which crosslinks with probe-bound RNAs or is quenched by bulk solvent. Upon total RNA isolation, RNA can undergo enrichment for Chem-CLIP or simultaneously processed for bulk RNAseq.

### Identification of putative RNA targets for NV1 considering enrichment and competition

High-quality cDNA libraries were generated (**Supp. Fig 1 (a)** and **(b)**) and pooled together for short-read sequencing (∼30-70 million reads/library). The majority of reads for all libraries were enriched in exonic regions of the transcriptome (**Supp. Fig 1 (c)**). In the analysis of the PAL-based Chem-CLIP samples, RNAs were defined as enriched if the relative abundance of the RNA following **NV1-PAL** treatment was at least two-fold greater than **B1-PAL** (p-value ≤ 0.05); for **NV1-PAL** 391 RNAs were enriched above **B1-PAL** (**Fig 3(a)**). RNAs were defined as competed if the relative abundance was decreased by at least two-fold in the presence of 20 µM **NV1** or **NV2** (p-value ≤ 0.05); 641 RNAs were competed by pre-treatment with **NV1** (**Fig 3(b)**).

**Figure 3:**
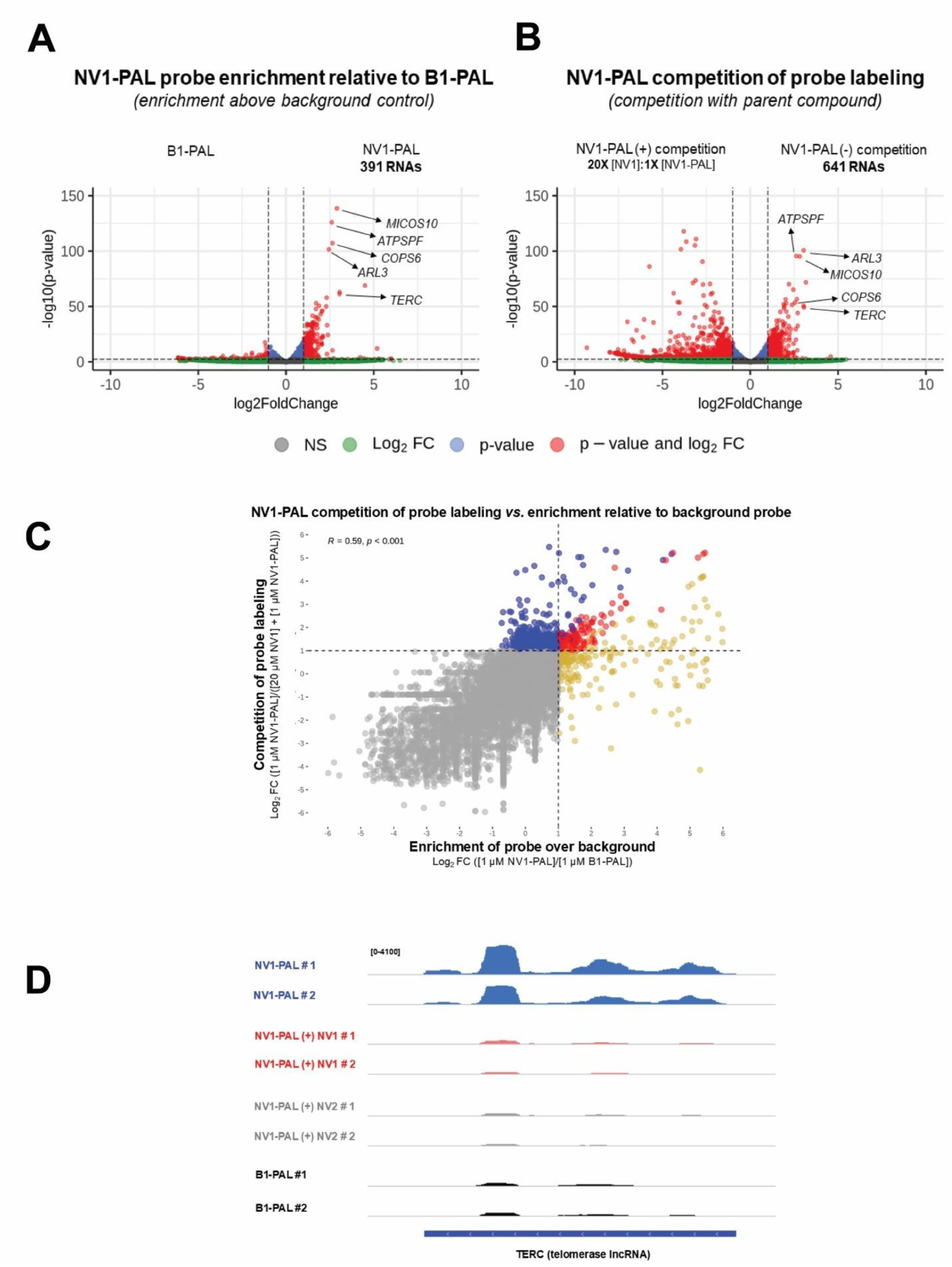
Unbiased cellular PAL-based Chem-CLIP profiling of NV1 reveals novel potential RNA targets. (A) Volcano plot of the relative transcript abundance comparing enrichment with **NV1-PAL** vs. **B1-PAL**. (B) Volcano plot of relative transcript abundance comparing enrichment with or without 20 µM **NV1**. Transcripts with differential enrichment were deemed hits if the fold-change ≥ 2 and p-value ≤ 0.05, as calculated using DESeq2^31^. (C) Scatterplot of differential enrichment (**NV1-PAL** vs **B1-PAL**) *vs.* competitive probe labeling (20 µM vs 0 µM **NV1**). Each circle denotes an individual gene (total transcripts/gene) with p-value ≤0.05. Gray points = fold change ≤2, Dark yellow = **NV1-PAL** enriched above **B1-PAL**, blue = **NV1** competed RNAs, and red = enriched and competed RNAs. Dotted lines denote fold-change cut-offs (FC ≥ 2). R^2^ coefficient = 0.59. (D) Read density of TERC across all conditions. Greater read density is observed for the **NV1-PAL** treatment alone. Genome track was visualized using Integrated Genomics Viewer (IGV) ^46^.

Based on established design criteria, RNAs statistically enriched beyond **B1-PAL** and whose probe labeling was competed by **NV1** were defined as hits; a total 161 RNAs were enriched and competed (**Fig. 3(c)** and **Suppl. Table 1**). Many of the top RNA hits were protein-coding mRNAs not previously shown to be modulated by **NV1**. We also observed a diverse set of putative RNA targets (**Supp. Fig 2(a)**) with a broad range of respective transcript abundances (**Supp. Fig 2(b)**). The strongest putative mRNA targets for **NV1** include ARL3, MICOS10, COPS6, and ATPSPF. An additional putative RNA target of **NV1** is the noncoding RNA TERC (telomerase lncRNA), the RNA component of the telomerase RNP complex which is localized to Cajal bodies and indispensable for telomere maintenance ^23, 24^. Read density from PAL-based Chem-CLIP experiments for TERC is shown in **Figure 3(d)**. In the **NV1-PAL** + **NV1** samples, TERC expression was found to increase relative to **NV1-PAL** alone, **NV1-PAL** + **NV2**, and **B1-PAL** treatments (not statistically significant); however, it is unclear whether this change in transcript abundance is a result of **NV1-PAL** engagement (**Supp. Fig 3**). For further reference the read density for GAPDH, a transcript that does not show enrichment or competition, is shown in **Supp. Fig 4.** For TERC, similar degrees of decreased read density resulting from competition of probe labeling with **NV1** and **NV2** are observed. Competitive binding in the presence of **NV2** suggests this putative binding interaction is unlikely to be functionally relevant in the context of splicing modulation. These data suggest a ligandable RNA/RNP binding pocket within TERC which has also been shown previously to bind a structurally unrelated riboswitch targeting metabolite, PreQ1, via PAL-based Chem-CLIP ^20^. With engagement observed with PreQ1 and **NV1,** we next wanted to ensure that TERC engagement was not an artifact of diazirine-based PAL labeling but rather **NV1** binding to a highly evolutionarily conserved ncRNA with important potential therapeutic relevance.

### Cellular Δ SHAPE-MaP profiling defines the CR4/5 domain of TERC as a likely binding site for **NV1**

For a PAL-independent cellular engagement assay, we turned to differential 2′-hydroxyl acylation reactivity coupled to mutational profiling (ΔSHAPE-MaP) ^25–27^ as a method to provide orthogonal evidence of binding by **NV1** in the same native cellular context as the PAL-based Chem-CLIP experiments. ΔSHAPE-MaP enables characterization of the small molecule binding site at single nucleotide resolution and provides insight into conformational changes in RNA secondary structure induced by compound engagement ^26, 28, 29^. Coupled to mutational profiling (MaP), chemically acylated nucleotides are miscoded during reverse transcription, and these mutational mismatches can be detected and read-out *via* NGS and read alignment.

To improve sequence coverage of TERC lncRNA, target TERC RNA was further enriched post cDNA generation *via* capture oligonucleotide probes. Sufficient sequencing depth was achieved to ensure 300X coverage of each residue of TERC (**Supp. Fig. 5(a)-(b)**). Significantly increased mutation frequencies were observed for only NAI probe treated samples (**Supp. Fig. 5(c)**). For the DMSO-treated cells, TERC exhibits distinct regions of high and low chemical probe reactivity indicating a highly structured RNA in cells (**Supp. Fig. 6(a)**). SHAPE-MaP performed in the presence of **NV1** produces a similar overall pattern of regions of differential high and low reactivity, suggesting that **NV1** does not globally alter the secondary structure of TERC RNA (**Fig. 4(a)**).

**Figure 4:**
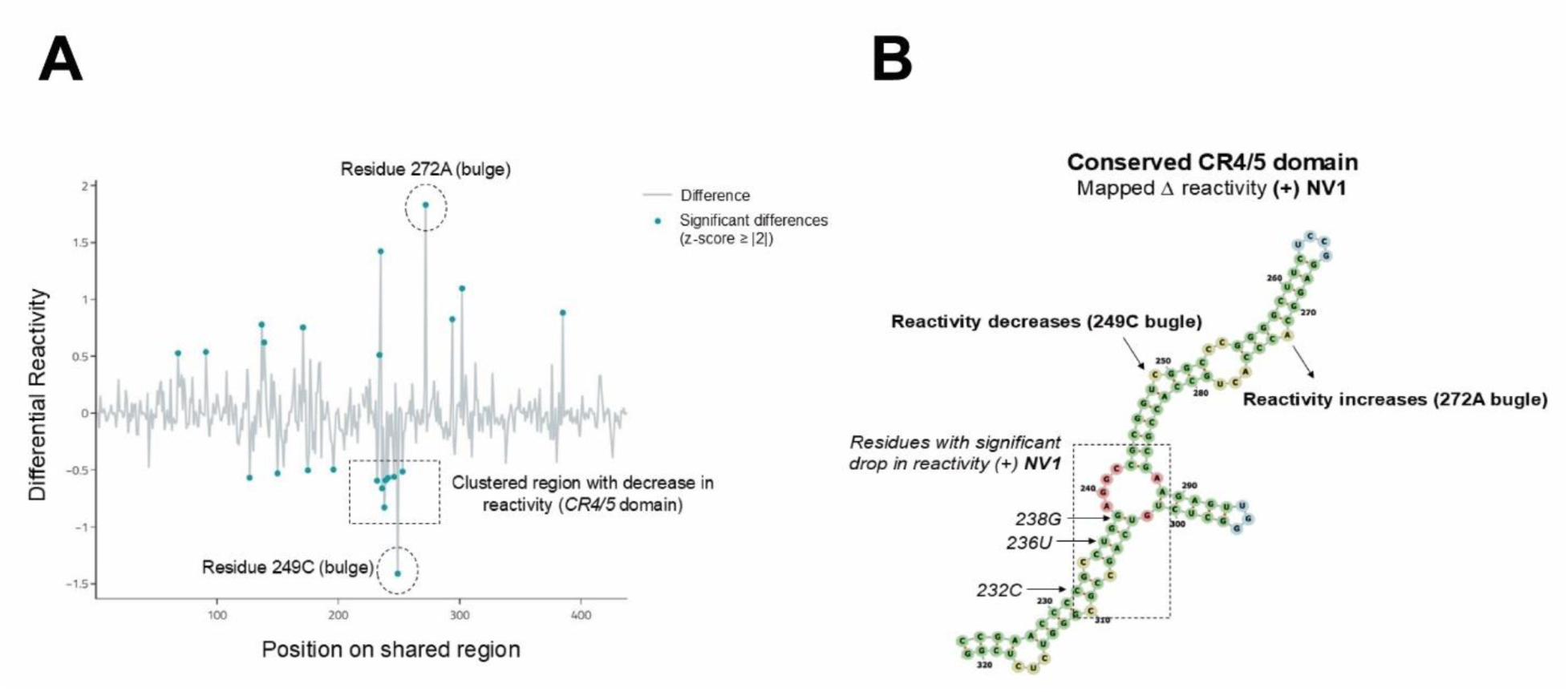
Cellular differential *SHAPE*-MaP of TERC lncRNA without and with NV1. (A) Single nucleotide resolution of differential reactivity for DMSO vs. 10 µM **NV1** treated samples. Blue dots indicate regions of significant differences in reactivity (z-score ≥ 2). Positive values indicate an increase in reactivity in the presence of **NV1**, while negative values indicate a decrease in reactivity. Noteworthy changes in reactivity are indicated by the boxed region within the conserved CR4/5 domain of TERC lncRNA. (B) Secondary structure within the CR4/5 domain of TERC lncRNA. Boxed region indicates site of significant clustered changes in chemical reactivity in the presence of compound. The bulged residue 249C shows the most significant reduction in reactivity and is likely important for NV1 binding. Secondary structures with SHAPE reactivity constraints were visualized using Forna ^44^. Nucleotides are colored based on structure (stems = green, multiloops/junctions = red, interior loops = yellow, hairpin loops = blue, 5’ and 3’ unpaired region = orange). All SHAPE-MaP experiments were performed with biological replicates (N=2) and include no NAI probe controls. Differential reactivities plotted 5’--> 3’ orientation of TERC lncRNA.

Applying ΔSHAPE, which compares chemical probe reactivity measurements between DMSO and **NV1**-treated samples, statistically significant changes in SHAPE reactivity were identified. **NV1**-treatment yielded both increases and decreases in SHAPE reactivity, indicating secondary structure rearrangements or differences in protein binding at the RNA binding interface upon **NV1** engagement. TERC RNA contains 5 distinct structural regions: Core pseudoknot, CR7, the H/ACA binding region, the hypervariable region, and the C4/5 domain ^24, 30^. Of these domains, the highly conserved C4/5 domain contained the greatest number and the strongest cluster of residues with decreased chemical probe reactivity in the presence of **NV1** (**Fig. (4a)**). We note that within the CR4/5 domain, C249 shows the strongest decrease in probe reactivity in the presence of **NV1** and may hint at this residue’s importance for **NV1** binding to TERC RNA (**Fig. 4 (b)** and **Supp. Fig 6(b)**). The observed changes in SHAPE-MAP reactivity provides the intended orthogonal validation that **NV1** likely binds directly with TERC RNA. Given, the number and magnitude of significant changes in SHAPE reactivity localized within the CR4/5 domain we propose that this domain is the likely site of compound engagement.

### SAR-dependent competition as a hit calling criteria enables identification of on-mechanism mRNAs

While further characterization of the **NV1**-TERC interaction *via* SHAPE reactivity profiling validates the PAL experiment itself, there were only a few mRNA expected targets ^1, 13^ of **NV1** that were also enriched and competed. While 161 RNAs were both enriched over **B1-PAL** and competed by **NV1**, many of these RNAs were ncRNAs and not expected on-mechanism mRNA splice targets (3’ nGA exon ending). In reviewing the two criteria for hit calling, we noticed that while known targets were able to meet the at least 2-fold competition criteria, they consistently failed to show greater than 2-fold enrichment over **B1-PAL**. Attempts to normalize or use other criteria to assess enrichment (differential peak calling, normalizing chem-CLIP read counts with bulk RNAseq, and transcript-dependent factor normalizations) repeatedly failed to show known targets as enriched (**Supp. Fig. 7**) and thus we could not find a method of analysis where the known targets would meet the designed two criteria for hit calling (enrichment and competition).

One of the advantages of multiplexing PAL-based Chem-CLIP samples with identically treated bulk RNASeq samples is that it allows for a direct comparison of enrichment through binding affinity versus transcript abundance. Comparing the relative transcript abundance following PAL-based Chem-CLIP with bulk RNASeq samples shows no covariation (**Fig. 5(a)**). This suggests that PAL-based affinity enrichment has an altered rank order of transcript abundance. In contrast, the relationship between transcript abundance in the competition samples for PAL-based Chem-CLIP versus bulk RNASeq shows a higher degree of positive covariation (R^2^=0.49) (**Fig. 5(b)**). This implies the competition samples (20 µM **NV1** + 1 µM **NV1-PAL**) are more like bulk RNASeq samples despite the presence of **NV1-PAL**. We interpret this to indicate that presence of competitor decreases PAL labeling, thus returning the relative abundance back towards their nonenriched states. Taken together, these observations indicate that competition, even without significant enrichment, may contain more relevant target engagement information than previously appreciated.

**Figure 5:**
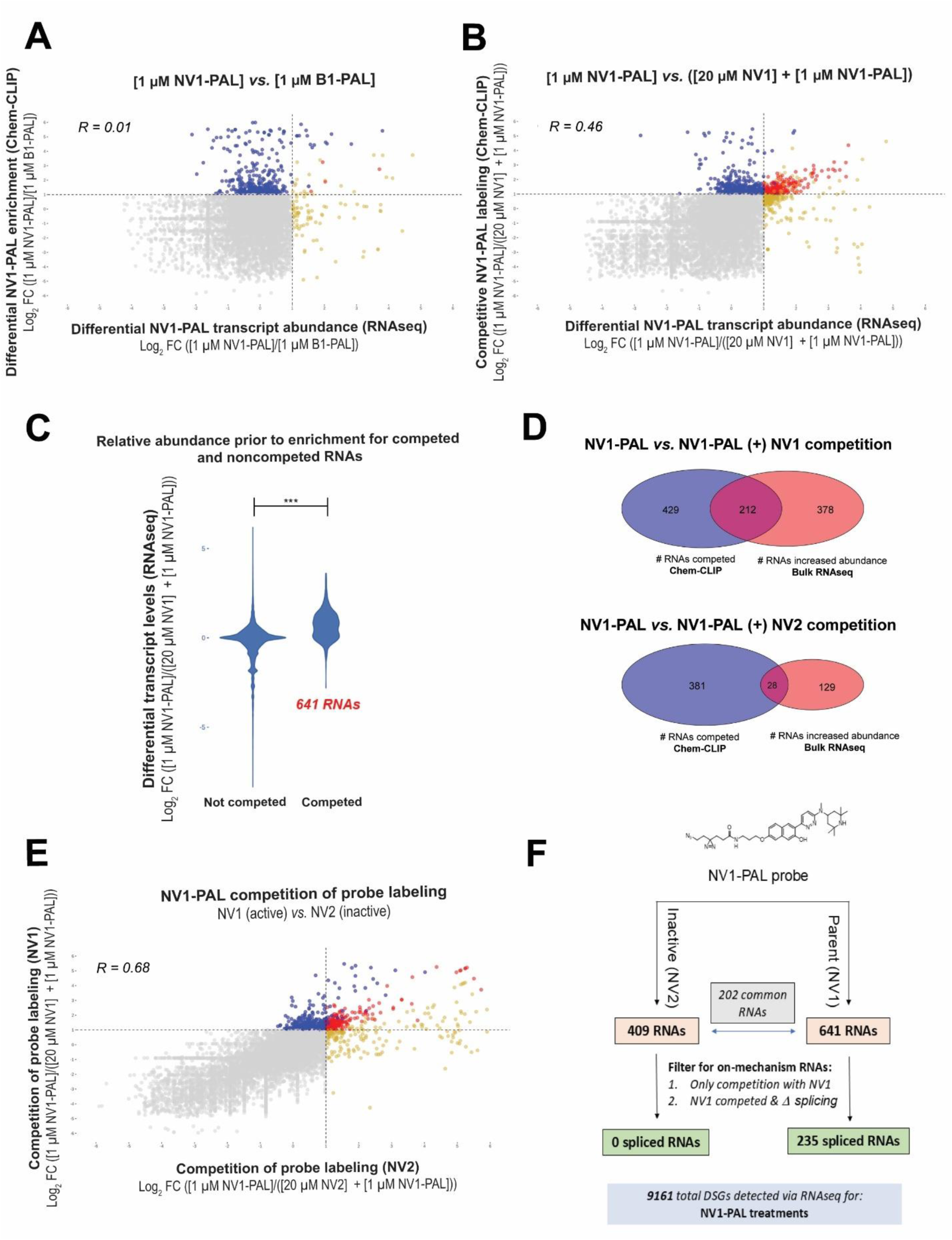
Refinement of hit calling strategy for small molecule RNA target ID. (A) Scatterplot of transcript abundance in the presence of **NV1-PAL** relative to **B1-PAL** (PAL-Based Chem-CLIP vs. bulk RNAseq). Only transcripts with p-value ≤0.05 are shown. Gray points = fold change ≤2, dark yellow = RNAs with increased relative abundance in the presence of **NV1-PAL** (RNAseq), blue = RNAs with increased relative abundance in the presence of **NV1-PAL** (PAL-based Chem-CLIP), and red = RNAs with increased relative abundance in the presence of **NV1-PAL** in both RNAseq and PAL-based Chem-CLIP. (B) Scatterplot of transcript abundance of 20 µM **NV1** + 1 µM **NV1-PAL** relative to 1 µM **NV1-PAL** (PAL-based chem-CLIP vs. bulk RNAseq). Only transcripts with p-value ≤0.05 are shown. Gray points = fold change ≤2, dark yellow = RNAs whose abundance is decreased by **NV1** treatment prior to PAL-based Chem-CLIP, blue = RNAs whose abundance is decreased by **NV1** treatment in PAL-based Chem-CLIP and red = RNAs whose abundance is decreased by **NV1** prior to and following PAL-based Chem-CLIP. (C) 2-way ANOVA analysis with Tukey’s honest significance test (HSD) was performed to assess statistical significance of differences in transcript levels prior to enrichment. Transcripts were binned as to whether they were or were not competed by **NV1** (PAL-based Chem-CLIP), x-axis. *** denotes p-value ≤ 0.001. RNAs whose PAL labeling was competed by **NV1** tend to also have lower transcript levels prior to enrichment. (D) Venn-diagrams of the overlap of PAL-based Chem-CLIP and bulk RNAseq hit calling. Lack of overlap suggests **NV1** competition in PAL-based Chem-CLIP hit calling is not driven by differences in transcript levels prior to enrichment. (E) Scatterplot of transcript abundance in the presence of 20 µM **NV1** vs. 20 µM **NV2**. Only transcripts with p-value ≤0.05 are shown. Gray points = fold change ≤2, dark yellow = RNAs specifically competed by **NV2**, blue = RNAs specifically competed by **NV1**, and red = RNAs competed by both **NV1** and **NV2**. (F) RNAs specifically competed by **NV1** were cross-referenced to differentially spliced genes (DSGs).

A potential cause for concern in using competition by **NV1**, irrespective of the magnitude of enrichment, as a hit calling criteria is that decreases in transcript abundance following **NV1** treatment may bias the data. It is well established that splicing modulation is often associated with a decrease in respective transcript levels ^8, 10, 13^. Indeed, we observe that RNAs competed by **NV1** have significantly lower transcript abundances relative to **NV1-PAL** treatment alone (**Fig. 5(c)**). However, statistically significant fold changes between competed transcripts (PAL-based Chem-CLIP) vs. transcripts with decreased abundance upon compound treatment (RNASeq) were distinctly different (**Fig. 5(d)** and **Supp. Fig 8**).

We note that while compound-competed transcripts were differentiated from transcripts with decrease abundance, there was some degree of overlap between these groups. To identify on-target splicing related hits, we filtered for transcripts whose enrichment was decreased by at least 2-fold in the presence of **NV1** and unaffected by **NV2** (inactive) (**Fig. 5(e)**). Recall that 20

µM **NV1** competed the enrichment of 641 transcripts. Of these, 202 RNAs were competed by both **NV1** and **NV2**, while 439 RNAs were selectively competed by **NV1** and not **NV2**.

Interestingly, among the 202 RNAs competed by **NV1** and **NV2**, many of these RNAs were also successfully enriched above **B1-PAL**. These include ARL3, MICOS10, COPS6, ATPSPF, and TERC (**Supp. Fig. 9**).

In terms of selective competition, 439 RNAs were exclusively competed by **NV1** and of these 235 were found to be differentially spliced in their corresponding bulk RNASeq samples (**Fig. 5(f)**, **Suppl. Table 2**). In contrast, 207 RNAs were competed by **NV2** alone and none of these were found to be differentially spliced in corresponding bulk RNASeq samples. The observation that only **NV1** competed RNAs were differentially spliced underscores that the known splicing specific mechanisms can be captured in the competition data despite any apparent sign of enrichment.

### SAR-dependent competition and not just competition alone is critical for hit calling

An important disclaimer about the competitor treated samples: we do not believe that most differentially spliced transcripts in the competition samples are the result of the U1snRNP molecular glue mechanism. Upon 1 µM **NV1-PAL** treatment, we observe 6116 unique differentially spliced genes relative to **B1-PAL** (background) using DEXseq ^32^. In contrast, the **NV1** competition samples (20 µM **NV1**+1 µM **NV1-PAL**) showed 9161 differentially spliced transcripts. This large difference in number of differentially spliced genes highlights a principal weakness of our study, namely using too large of an excess of competitor ligand. Hence, our focus on only RNAs selectively competed by **NV1** but not **NV2** as potential **NV1** targets.

### Novel putative target identified from unbiased profiling

One of the additional value propositions of unbiased profiling is the access to novel target engagement data. While assessing expected targets of **NV1**, we noticed that GATA3 was competed by **NV1** and differentially spliced (**Fig 6(a)**), suggesting possible target engagement. GATA3 is a transcription factor implicated in breast cancer prognosis ^33^. GATA3 has a 3’ AGA exon ending containing transcript isoform. Interestingly, **NV1** treatment was associated with a decrease of exon inclusion of the 3’ AGA exon within GATA3, as well as a decrease in inclusion of other exons within the transcript. These splicing changes were also associated with a significant reduction in total GATA3 transcript abundance in the (20 µM **NV1** + 1 µM **NV1-PAL**)-treated samples (**Fig 6(b)**). Collectively, these data suggest **NV1** engagement inhibits exon inclusion and ultimately leads to decrease in GATA3 transcript abundance. Of the 235 RNAs competed and differentially spliced in the presence of **NV1**, 10 RNAs contain the expected 3’ nGA exon motif (**Fig. 6(c)**). It is important to recall that these experiments were performed with a 4 hr treatment duration and likely fail to capture the total 3’ nGA exons that were likely compound-induced in other reported experiments with longer treatment times (typically > 24hrs).

**Figure 6:**
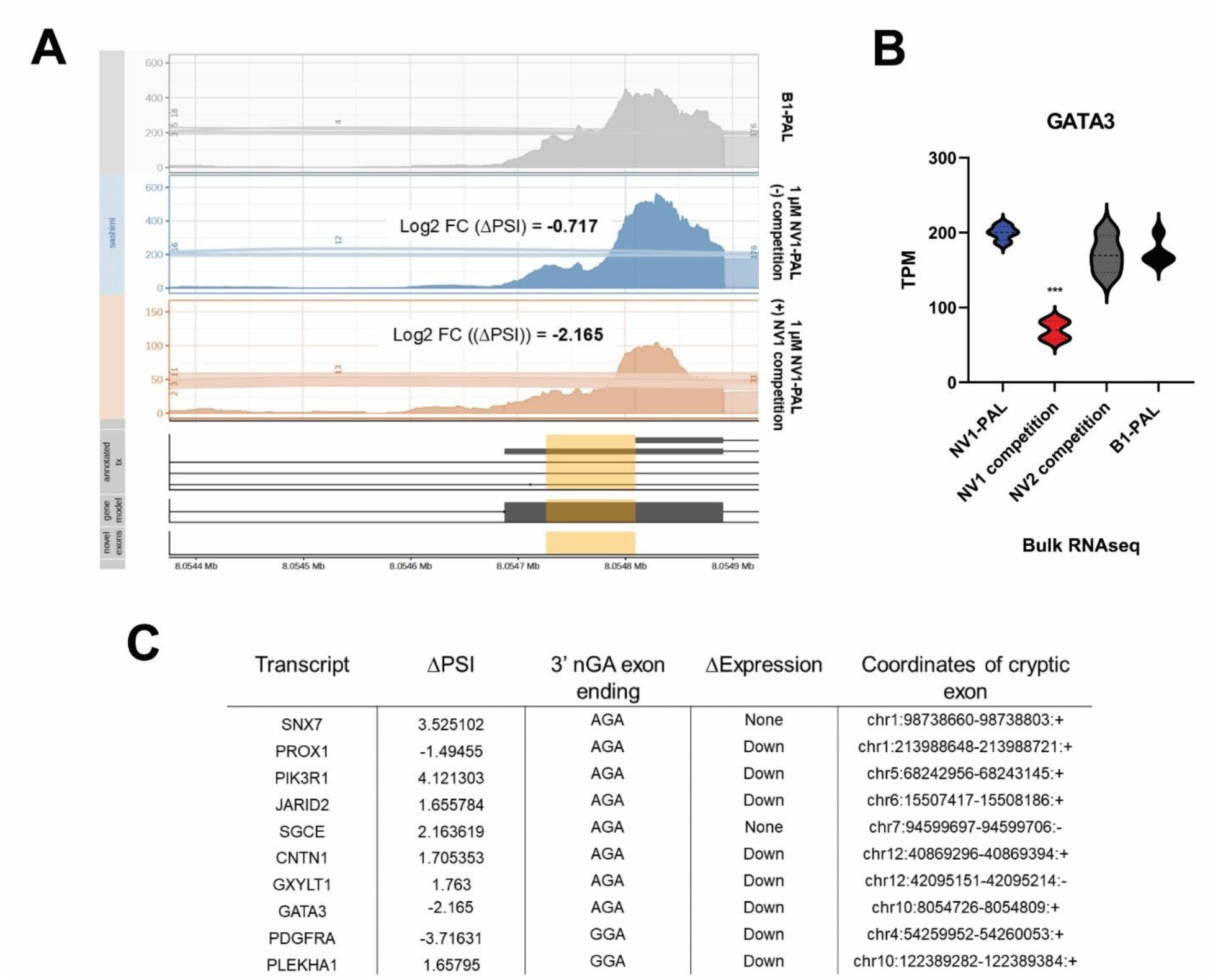
Targetability of GATA3 revealed through PAL-based Chem-CLIP. (A) Sashimi splicing plot at 3’ AGA exon site at *GATA3* locus. Highlighted region denotes cryptic splice site. Significant decrease of exon inclusion observed for both **NV1-PAL** and **NV1-PAL** with 20 µM **NV1**. Δ Percent spliced in (PSI) is relative to differential splicing observed with **B1-PAL** treatment. A positive PSI indicates increase in splicing inclusion, while a negative value indicates a decrease of inclusion. (B) TPM values of total GATA3 transcripts across compound treatments. *** denotes a p-value ≤ 0.001 for differences in TPM values (transcriptome-wide student T-test). (C) Table of 10 RNAs with 3’ nGA endings that are competed by **NV1** and differentially spliced.

## Conclusion

Previous RNA target identification studies using covalent chemical probes ^18–20, 22, 34^ have demonstrated feasibility of covalent probe-mediated RNA enrichment coupled to NGS. These pioneering studies leveraged exogenous expression of high-affinity aptamers ^19, 20^ or were performed in cellular lysates, establishing critical proof-of-concept and forming foundational guidelines for transcriptome-wide profiling of RNA-small molecule interactions. Building on these prior studies, our aim was to profile the RNA targeting landscape of a class of clinically relevant splicing modulators under endogenous cellular conditions.

Using conventional hit calling criteria, we identified the lncRNA TERC as a previously unknown target of **NV1** and **NV2**. Further characterizing the **NV1**-TERC interaction with Δ SHAPE-MaP highlighted a defined region within the CR4/5 domain of TERC showing decreased chemical reactivity. Further work is needed to better define the binding of **NV1** with TERC at a molecular level. While we propose initially focusing such efforts on the CR4/5 domain, we remain open to the possibility that **NV1**-TERC engagement is protein-mediated or via a binary RNA-interacting mechanism analogous to BIBR1532, a previously described TERC inhibitor ^35^.

To enable the advancement of translating PAL-based Chem-CLIP into an endogenous cellular system, we leveraged knowledge starting from a well-characterized splicing tool compound to determine experimental parameters and target hit calling criteria. The extensive characterization of U1snRNP-dependent molecular glues enabled us to pinpoint that lack of enrichment as the failing criteria in our study. Also, we leveraged this information to focus on competition, even in absence of enrichment, as a meaningful way to interpret the data.

While we demonstrate that early binding events for **NV1** are consistent with expected on-mechanism mRNA hit calling from expression profiling studies, we also capture engagement with diverse RNA biotypes that may be a product of compound cellular distribution or suggest engagement with structured RNA/RNP active sites. The latter would indicate potential druggable RNAs that could be developed into functional modulators through traditional medicinal chemistry or by imparting functionality into the compounds; for example, RIBOTACs or covalent RNA binders^36–38^.

## Methods

### General details for photoaffinity probe synthesis and characterization

Detailed chemical synthesis schemes with corresponding characterization of chemical photoaffinity probe can be found in the supplemental information. General operating procedures of LC/MS, high resolution mass spectral analysis, and HNMR can be found below.

LC/MS Analysis:

LC/MS analysis of mass and purity was carried out on Waters ACQUITY UPLC BEH C18, 130Å,

1.7 µm, 2.1 mm X 50 mm. Solvent A: water +0.1% formic acid. Solvent B: acetonitrile +0.1% Formic Acid. Average column temperature: 50.0 °C,. 1 µLA 2.0-μL aliquot of an analytical solution was injected and eluted at flow rate of 250 μL/min with a 5.2 min combination of both isocratic and linear gradient elution using mobile phase varying from 2-98% Solvent B.

High Resolution Mass Spectral Analysis:

LC/MS analysis of accurate mass measurements was carried out on the Waters Acquity CM system. Solvent A: water + 0.05% TFA. Solvent B: acetonitrile + 0.05% TFA, column type: ACQUITY UPLC® CSH™ C18 1.7μm X 50 mm, A aliquot of an analytical solution was injected and eluted at flow rate of 1 mL/min with a 2.2 min combination of both isocratic and linear gradient elution using mobile phase varying from 2-98% Solvent B average column temperature: 50.0 °C.

HNMR system:

Avance III 400 NMR spectrometer, 1H-frequency: 400.13 MHz

### Cell culture conditions

The adherent human neuroblastoma bone marrow biopsy-derived cell line, SH-SY5Y cells (Sigma catalog: 86012802), were cultured in DMEM/F-12 (Gibco, cat: 11320033) supplemented with 10% FBS and 1% penicillin/streptomycin. All cells were cultured at 37°C in 5% CO2 in humidified incubators and were free from mycoplasma (MycoAlert Detection Kit, Lonza).

### Bioactivity characterization of diazirine-based PAL chemical probes

Bioactivity of chemical probes relative to parent compound was tested using the *SMN2* mini-gene luciferase reporter (SH-SY5Y cells). 1.2 × 10^4^ cells were seeded into individual wells of two separate 384-well plates for activity and cytotoxicity assays and allowed to recover at 37°C with 5% CO2 overnight. The following day, serial dilution of chemical probe or parent compound was added, then following 24 hr incubation, the luciferase and cytotoxicity assays readings were performed. Luciferase expression and cytotoxicity was assessed using the Promega Bright-Glo luciferase assay and CellTiter-Glo luminescent cell viability assay, respectively. All assays were performed triplicate.

### Cellular treatment for PAL-based Chem-CLIP workflow

For the PAL-based Chem-CLIP workflow, all treatment conditions were performed in triplicate. 1.2×10^7^ SH-SY5Y cells were seeded into individual 10cm^2^ cell culture plates using FluoroBrite DMEM media (Gibco, cat: A1896701) supplemented with 1% FBS and allowed to incubate at 37°C with 5% CO2 overnight. Cells were then treated with either DMSO or 20 µM competitor compound for 1 hr at 37°C with 5% CO2. Next, cells were treated with either DMSO or 1 µM PAL probe for 3 hrs at 37°C with 5% CO2. Prior to photocrosslinking, cells were washed once with DPBS (Gibco, cat: 14190144), after which 5 mL of cold DPBS was added to each plate. Photocrosslinking was performed for 20 min at 4°C with 365 nm wavelength light (UVP Blak-Ray XX-40BLB UV bench lamp, 40 Watt). Cells are then maintained at 4°C and harvested *via* scraping. The detached cells and media were pelleted (300×*g* for 5 minutes at 4°C). The supernatant was aspirated and the cell pellets flash frozen and stored at –80°C until further processing.

### RNA extraction for PAL-based Chem-CLIP workflow

Cells are lysed and RNA is extracted following the Qiagen RNeasy Plus (cat: 74134) following manufacturer’s protocol. Cell pellets were resuspended in 600 µL of RLT lysis buffer and homogenized using the Qiagen QIAshredder (cat: 79656). 600 µL of 70% ethanol was added to the flow-through of the QIAshredder column and transferred to a gDNA column provided by the RNeasy kit. The remainder of the RNeasy protocol is followed for total RNA isolation including on-column DNase digestion using the Qiagen DNase set (cat: 79254). Quality and quantification of purified RNA is evaluated on a Nanodrop prior to further processing.

### Copper-free click chemistry for PAL-based Chem-CLIP workflow

Copper-free click reactions were performed in nuclease-free water with 10 µg of total RNA, 10 µM final concentration of DBCO-PEG4-desthiobiotin (Sigma Aldrich cat: 760749) with 1 unit/µL final concentration of SUPERase·In™ RNase Inhibitor (Invitrogen cat: AM2694). The click-reaction was allowed to proceed at room temperature for 2 hrs with gentle agitation. Following completion of the copper-free click chemistry reaction, total RNA was purified from other reaction components using a Zymo RNA spin column (Zymo Research, cat = C1003-50). RNA was eluted from the columns with 22 µL of nuclease-free water. Biotinylated-RNA was stored at –20°C until ready for processing.

### Enrichment of biotinylated RNAs for PAL-based Chem-CLIP workflow

Magnetic streptavidin beads (Invitrogen Dynabeads MyOneStreptavidin C1 (cat# 65002)) were blocked in batch at room temperature in a blocking buffer (0.03% BSA, 10 µM yeast tRNA (Invitrogen, cat # AM7120G)) with end-over-end agitation. Ensure sufficient beads to allow for 20 µL of beads/sample during enrichment step. Following blocking, beads are washed three times with wash buffer (50 mM Tris-HCl (pH 7.4), 100 mM NaCl, 10 mM EDTA (pH 8.0), 0.2% Tween-20, 1 unit/µL SUPERase·In™ RNase Inhibitor), then resuspended to make a slurry and aliquoted to individual tubes to provide 20 µL of beads/sample. When ready to proceed to final enrichment step, the beads were separated from the wash buffer using a magnetic stand, then spun at 200xg for 1 min and any residual wash buffer was removed by aspiration.

For enrichment with streptavidin, 20 µL of Zymo purified RNA from the click reaction was added to individual aliquots of beads. Incubation of streptavidin beads with RNA proceeded for 2 hrs at room temperature with end-over-end agitation. Following this incubation, the supernatant was removed and beads were washed three times with 200 µL wash buffer, then with 200 µL of nuclease-free water to remove residual wash buffer which contains EDTA and may impair reverse transcription efficiency.

### On-bead reverse transcription and cDNA generation for PAL-based Chem-CLIP workflow

Following streptavidin-mediated enrichment of photocrosslinked RNAs, reverse transcription is performed directly on beads. The SMARTer Stranded Total RNA-seq kit v3 – Pico input (Takara, cat: 634486) was used with some modification from the manufacturer’s protocol. Magnetic streptavidin beads with bound biotinylated target RNAs were resuspended in 10 µL of nuclease-free water.

To fragment probe-labeled RNAs, 8 µL of resuspended RNA-bead mixture was added to individual wells of a PCR plate containing 1 µLSMART Pico Oligos Mix v3, 4 µL5X First-Stranded Buffer and incubated at 94C for 3.5 min, then immediately placed on ice to halt further fragmentation. The manufacturer’s protocol for reverse transcription reaction was followed. It is worth noting to take extra precautions to homogenize the reverse transcription reactions as the solution is viscous.

Following completion of the first strand synthesis reaction, 20 µL of each reaction mixture (beads and solution) was transferred into individual wells of a PCR plate. RNase H cleavage reactions were initiated with the addition of 5 µL10X RNase H reaction buffer, 24 µL nuclease-free water, and 1 µL RNase H (New England Biosciences, cat #M0523). RNase H cleavage reaction proceeded at 37°C for 20 minutes with gentle agitation. RNase H cleavage reaction was quenched with addition 1 µL0.5 M EDTA (pH 8.0).

Magnetic streptavidin beads were removed from the reaction solution with aid of magnetic stand. First strand synthesized cDNAs were further purified using RNAclean XP beads (Beckman Coulter, cat# A63987) with the addition of 100 µL of RNAclean XP beads to each reaction mixture, incubated at room temp for 5 min. Supernatant was removed with aid of magnetic stand. RNAclean XP beads were washed twice with 100 µL70% ethanol. Carefully pipette residual ethanol from samples and allow beads to air dry for 10 minutes. Single-stranded cDNA was eluted from beads with 22 µL nuclease-free water and proceed to first PCR amplification step in library generation per manufacturer protocol including ribodepletion until final PCR amplification step. The number of final PCR cycles may require optimization depending on crosslinking, enrichment efficiency, and RNA quality. Our final libraries were generated with 14 PCR cycles for PAL-based Chem-CLIP cDNA libraries. Final library quality was assessed using a D1000 TapeStation system (Agilent, cat = 5067-5582).

### Reverse transcription and cDNA generation for bulk RNAseq

Wildtype SH-SY5Y cells were treated identically and in parallel with cellular treatment for PAL-based Chem-CLIP workflow. RNA was harvested identically and in parallel with RNA extraction for PAL-based Chem-CLIP workflow. Reverse transcription and cDNA generation were performed as described for On-bead reverse transcription and cDNA generation for PAL-based Chem-CLIP workflow, except that 10 ng of total RNA input was used in place of Magnetic streptavidin beads bound to biotinylated target RNAs. Also, during the final PCR amplification step, 10 PCR cycles were performed.

### NGS sequencing parameters for PAL-based chem-CLIP and bulk RNAseq

Libraries were diluted to 5 nM in TE Buffer (pH 8.0), pooled, denatured, and loaded on the Illumina NovaSeq 6000 instrument (Part number 20012850) for sequencing following the manufacturer’s instructions. The PAL-based Chem-CLIP libraries were loaded at a final concentration of 300 pM with a 6% PhiX spike in to increase sequence diversity. The bulk RNAseq libraries were loaded at a final concentration of 100 pM with a 1% PhiX spike, per manufacturer’s recommendation. Each library set was independently loaded and sequenced using the NovaSeq 6000 SP Reagent Kit v1.5 (Part number 20027466) per the manufacturer’s instructions. The PAL-based Chem-CLIP libraries were sequenced 53 base pair paired-end with 8 base dual indexes and trimmed to 50 base pairs. The bulk RNAseq libraries were sequenced 51 base pair (paired-end) with 10 base dual indexes and trimmed to 50 base pairs. The sequence intensity files were generated using the Illumina Real Time Analysis software (RTA3). An internal data processing pipeline was used to demultiplex, create, extract and trim the FASTQ files using Illumina’s bcl2fastq software (v2.20). An average of 30 million reads and 40 million reads per sample were obtained for the PAL-based Chem-CLIP and bulk RNAseq libraries, respectively, with a minimum of 23 million reads per sample.

### Analysis of PAL-based Chem-CLIP and RNAseq datasets

A previously reported exon quantification pipeline (version 2.5)^39^ with STAR (version 2.7.3a)^40^ was used to align the reads against the human genome reference from the latest build (GRCh38). Differential gene expression and differential PAL-based Chem-CLIP enrichment were determined using DESeq2 (v1.38.3)^31^. RNAs were defined to be differentially expressed or enriched, respectively, if |log_2_FC| ≥ 2 and p-value ≤ 0.05 relative to **B1-PAL**. Alternative strategies for normalization of PAL-based Chem-CLIP read counts apply transcript-dependent normalization, where enriched read counts are matched with a bulk RNAseq (transcript abundance) read count matrix. Normalization matrices are then set to a row-wise geometric mean ∼1. Sample code can be found in **Supp Fig 7**.

Splicing analysis of bulk RNAseq samples was performed using the DEXseq splicing analysis pipeline, an algorithm for detecting differentially spliced exons by surveying junction reads^32^. Changes in splicing and gene expression were correlated to PAL-based Chem-CLIP to understand if compound engagement leads to a functional response.

### RT-qPCR to assess splicing efficiency of PAL probes & qPCR read-out of PAL-enriched cDNA libraries

SMN2 mini-gene expressing SH-SY5Y cells were plated into individual wells of 6-well plate (1×10e6 cells/well) and allowed to recover at 37°C with 5% CO2 overnight. Cells were treated 1 µM for the indicated time points. Cells are harvested, and total RNA is isolated as described in *RNA extraction for PAL-based Chem-CLIP workflow*. Reverse transcription is performed following the manufacturer’s protocol for SuperScript II (Invitrogen, cat# 18064-014) and qPCR was performed using the 2X SYBR Green qPCR master mix (Invitrogen, cat = A46109) using standard cycling conditions (50°C for 2 minutes (min), 95°C for 2 min followed by 40 cycles of 95°C for 15 seconds (s) and 60°C for 60 seconds). Primer sequences used to assess splicing kinetics was 5’-GAAGGAAGGTGCTCACATTCCT-3’ (SMN2 exon7, Forward) and 5’-GGCGCCGGGCCTTTCTTTATG-3’ (Luciferase, Reverse).

PAL-enriched cDNA libraries can also be generated using the SuperScript II protocol, performing the RT reaction directly on the magnetic streptavidin beads. Following first-strand cDNA synthesis, a subsequent RNase H reaction and cDNA clean-up is performed as detailed in the chem-CLIP protocol to release single-stranded cDNA from the photocrosslinked RNA templates. The final cDNA can then be used as input for qPCR using the 2X SYBR Green qPCR master mix. This procedure is how we tested photocrosslinking to target RNAs in cells in **Fig. 1(e)**).

*Cellular SHAPE-MaP cDNA library generation:*

SH-SY5Y cells were plated in 10cm^2^ cell culture plates with DMEM/F-12 antibiotic-free media supplemented with 10% FBS. The following day, cells were treated with DMSO or 10 µM **NV1** for four hours. Following treatment duration, cells were harvested (cell scraper), spun down (300x*g*), resuspended in 950 µL of room temperature PBS. 50 µL of DMSO (background mutation control) or 50 µL of 2M NAI (EMD Millipore; Cat# 03-310) were added to the resuspended cell mixture for 15 minutes at 37°C with gentle agitation. DMSO samples were processed initially to minimize over acylation. Following NAI treatment, samples were spun down in 4C centrifuge (1000xg) for 5 minutes, and supernatant was aspirated without disturbing the cellular pellet. Pellets were washed and resuspended in chilled PBS. Samples were spun down again, and PBS was aspirated without disturbing pellets. Cellular pellets were flash frozen and stored at –80°C. Total RNA was isolated as described in *RNA extraction for PAL-based Chem-CLIP workflow*, except that the RNA was eluted in 50 µL nuclease-free water. Final sample quality was assessed on an RNA TapeStation and concentration/purity was assessed with the Nanodrop.

10 μg of total RNA isolated from each SHAPE-treated sample was subjected to rRNA depletion, and 100 ng of rRNA-depleted RNA samples were fragmented and subsequently phosphorylated in preparation for ligation. After phosphorylation, barcoded RNA adapters were ligated to the 3′ end of the fragmented RNA. RNA was reverse transcribed in the presence of MnCl_2_ and betaine as previously described ^25^ to induce mutations in the cDNA followed by ssDNA adapter ligation, quantitative PCR, and PCR amplification to generate sequencing libraries. 500 ng of each final library was pooled, and probe hybridization was performed followed by biotin-based enrichment of target sequences. 5′-biotinylated enrichment probes were designed to tile the target sequence using the FASTA sequence for ENSG00000270141, and a custom panel was created by mixing equimolar concentrations of each probe. Custom probes were designed and purchased from Integrated DNA Technologies. Following hybridization to target TERC lncRNA, biotinylated probes hybridized to the target complex were enriched *via* magnetic streptavidin beads. Beads were washed using the wash buffer (50 mM Tris-HCl (pH 7.4), 100 mM NaCl, 10 mM EDTA (pH 8.0), 0.2% Tween-20, 1 unit/µL SUPERase·In™ RNase Inhibitor). Post-enrichment amplification was performed on the purified target sequences and quantified on an Agilent 4200 TapeStation. The enriched cDNA pool was sequenced on a NextSeq 1000/2000.

### Cellular SHAPE-MaP analysis

Reads were barcode trimmed with cutadapt 1.14 and mapped to a custom reference genome generated using the primary sequence of TERC lncRNA. There was incomplete coverage using the sequence for ENSG00000270141, thus the primary sequence for telomerase-vert (RF00024) was used (ENSG00000277925) to generate the reference genome. Star aligner version 2.4.0i was used for reference alignment, and duplicate reads were filtered using umItools 0.5.0. Downstream analysis was performed as previously described ^25^ using the ShapeMapper 2.0 pipeline ^25, 41, 42^.

The output of SHAPE reactivity scores were then used as constraints for secondary structure prediction using RNAstructure ^43^. The secondary structure generated using SHAPE reactivities was exported to the forna visualization software ^44^ to compare secondary structures of DMSO with **NV1** treated samples.

### Curation of differentially expressed exons with 3’ nGA exon endings for quantification of 3’ nGA exon ending enrichment in NV1 probe target profiles

Starting with the splicing output from DEXseq ^32^ to compile splicing junctions of NV1 treated cells from the bulk RNAseq analysis, bed files were created using the exon boundaries. The primary sequence of these respective boundaries from the GRCh38 reference genome build were extracted using bedtools^45^. Exons with 3’ nGA endings were then further extracted and cross-referenced back to the NV1-PAL RNA target profiles to identify putative RNA interactors with possible 3’ nGA ending cryptic exons.

## Data Availability

PAL-based Chem-CLIP, bulk RNAseq, and SHAPE-MaP raw sequencing files have been deposited on SRA with the associated BioProject (####). Custom analysis scripts for data processing and quality control can be provided upon request. Data supporting the findings of this study not found within the main text of the paper can be found in the Supplementary Information files.

## Acknowledgements

The authors would like to thank Razvan Nutiu for his support in earliest phases of these studies and Steve Canham for engaging discussions and review of the manuscript. We also thank Eclipse Bioinnovations for their help with SHAPE-MaP sample preparation and analysis. R.S. was supported by the Novartis Biomedical Research Postdoctoral Office.

