## Supplemental Figures for "Photoaffinity enabled transcriptome-wide identification of splice modulating small molecule-RNA binding events in native cells"

Novartis Biomedical Research, Discovery Sciences, Cambridge, MA, USA

**Content:** Supplementary Figures **S1-10** and chemical probe synthesis/characterization (**Section II**). Excel tables provided with detailed information to reproduce plots in main text:

1. Table of 161 RNA hits enriched above **B1-PAL** & competed by **NV1**
2. Table of 235 RNAs competed by (**NV1** & not **NV2**) & differentially spliced
3. Differentially spliced genes (DSGs) for all **NV1-PAL** treated samples compared to **B1-PAL** (background) treatment (DEXseq [1] output)
4. Differential enrichment (Chem-CLIP) and expression (RNAseq) log2FCs & p-values provided for all sample contrasts
5. TPM values for Chem-CLIP & RNAseq samples (all biological replicates)
6. SHAPE reactivity values for DMSO- & **NV1**-treated samples from ΔSHAPE-MaP experiments

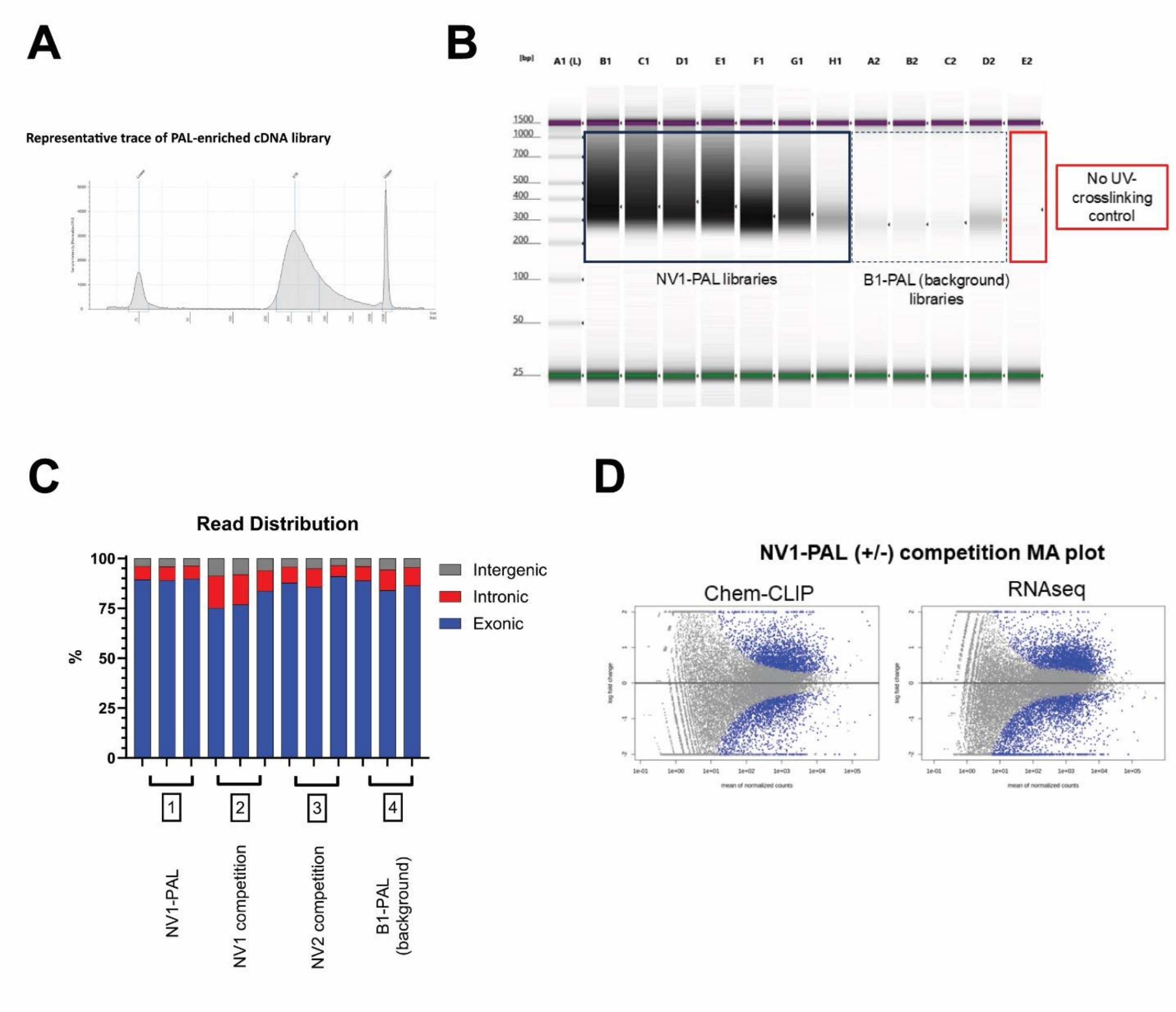


**Supplemental Figure 1. Quality control assessment of PAL-based Chem-CLIP libraries.** (A) Representative chromatogram from TapeStation trace of final PAL-based Chem-CLIP cDNA library prior to pooling and sequencing (B) PCR amplified product of final PAL-based Chem-CLIP cDNA libraries visualized on TapeStation gel. Average fragment sizes are 300-500 bp per library with final product concentration ranging from 14-150 nM. Increased final product observed for **NV1-PAL** enriched libraries relative to empty PAL control. Little to no product observed in no UV irradiation or DMSO controls (C) Read distribution of PAL-based chem-CLIP libraries analyzed in this study. (D) Minus-Average (MA) plots (normalized counts *vs.* $\log_2$ fold changes) of respective RNA transcripts to evaluate magnitude of fold changes relative to mean changes in enrichment relative to **NV1-PAL** with 20 µM **NV1-PAL** samples. MA plots were visualized to compare differential profiles of parallel processed PAL-based Chem-CLIP *vs.* RNAseq libraries for **NV1-PAL** treated samples without or with 20 µM **NV1-PAL**.

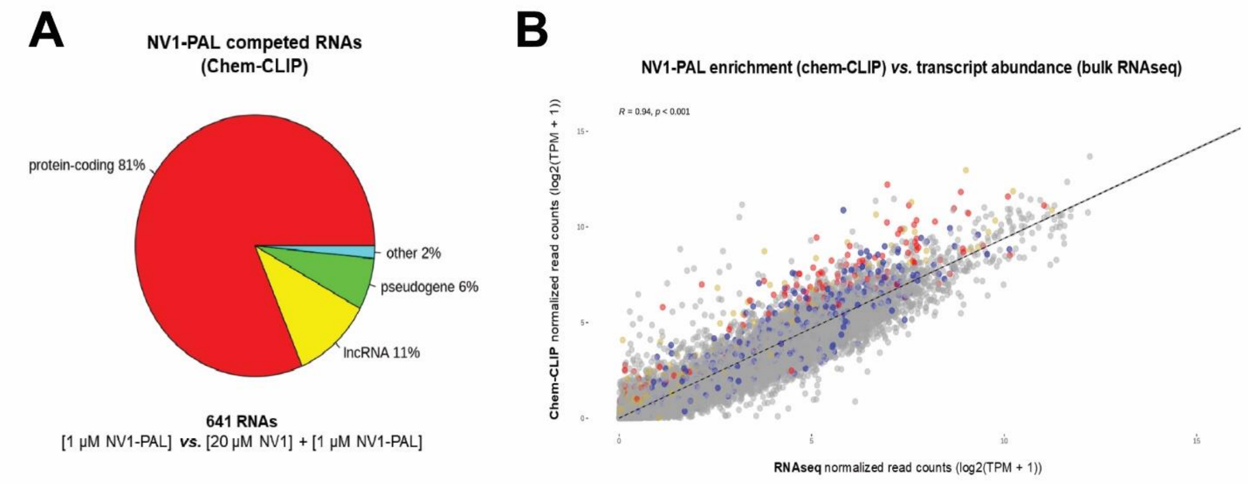


**C**

| Δ transcript expression (DEGs) | Δ expression of NV1 competed RNAs |
|---|---|
| Up-regulated | 1 |
| Down-regulated | 212 (35%) |
| No change | 428 (66%) |

**Supplemental Figure 2. Descriptive RNA hit profiles of NV1-PAL probe**. (A) RNA biotype distribution of **NV1** competed RNAs. (B) Scatterplot of **NV1-PAL** normalized read counts from PAL-based chem-CLIP experiments *vs.* the normalized read-counts from the parallel RNAseq experiment. $R^2$=0.94, suggesting strong association between degree of PAL labeling and total transcript abundance. Grey points = no significance, dark yellow = RNAs enriched by **NV1-PAL** over **B1-PAL**, blue = RNAs competed by **NV1**, red = enriched and competed RNAs. Significance = $\log_2$FC ≥ 1.0 and p-values ≤ 0.05. (C) Distribution of changes in expression for **NV1-PAL** competed RNAs.

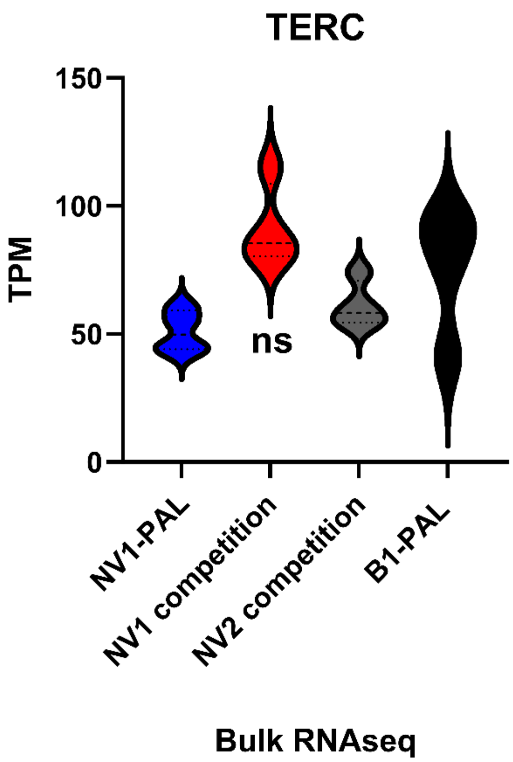


**Supplemental Figure 3. TPM values (RNAseq) of TERC lncRNA across PAL treatment conditions.** (A) TPM values of total TERC transcripts across compound treatments. ‘ns’ denotes a non-significant p-value for differences in TPM values (transcriptome-wide student T-test).

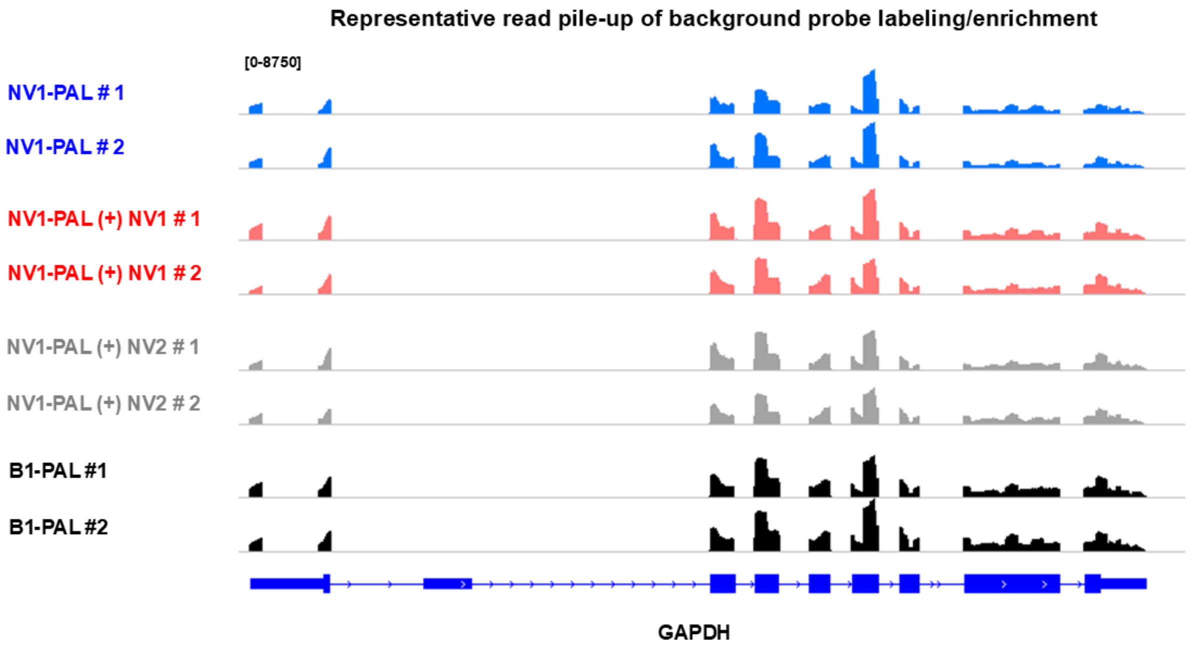


**Supplemental Figure 4. Read density of *GAPDH* locus across all treatment conditions.** Comparable read counts across all treatment conditions at GAPDH, demonstrating background signal in this assay.

**A**

| DMSO (v/v) or NV1 | NAI or DMSO (v/v) | Initial reads | % Pass trim | % Uniquely aligned | % PCR duplicates | Final reads | % Covered bases | Mean coverage |
|---|---|---|---|---|---|---|---|---|
| DMSO | DMSO | 18,530,655 | 99.84% | 0.38% | 42.07% | 40,970 | 100.00% | 5,655 |
| DMSO | DMSO | 20,218,636 | 99.87% | 0.44% | 40.71% | 52,508 | 100.00% | 7,173 |
| DMSO | NAI | 18,979,226 | 99.83% | 0.34% | 43.91% | 36,611 | 100.00% | 4,898 |
| DMSO | NAI | 22,471,126 | 99.85% | 0.31% | 44.01% | 38,529 | 100.00% | 5,257 |
| NV1 | DMSO | 20,212,955 | 99.84% | 0.50% | 42.54% | 57,504 | 100.00% | 7,635 |
| NV1 | DMSO | 20,133,344 | 99.81% | 0.42% | 44.49% | 46,909 | 100.00% | 6,274 |
| NV1 | NAI | 22,037,658 | 99.84% | 0.33% | 45.98% | 38,901 | 100.00% | 5,359 |
| NV1 | NAI | 21,659,504 | 99.83% | 0.28% | 43.79% | 33,885 | 100.00% | 4,654 |

**B**

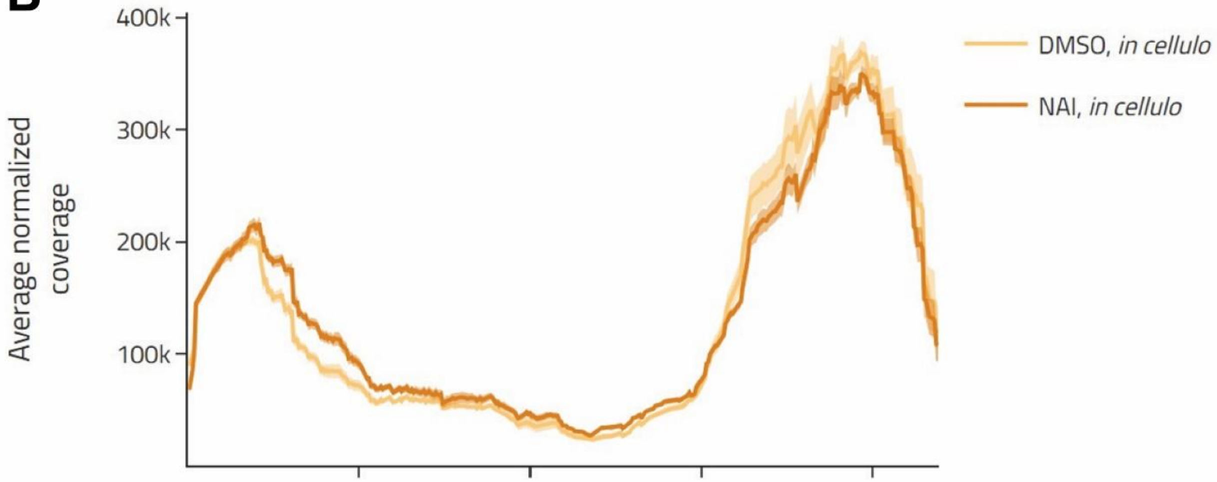


**C**

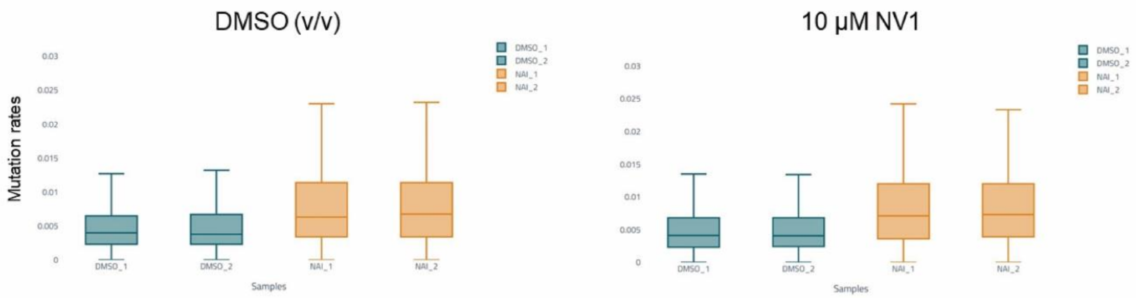

**Supplemental Figure 5. Cellular SHAPE-MaP library quality control assessment.** (A) Summary statistics of sequencing & read alignment at TERC locus for all SHAPE-MaP samples. (B) Normalized read coverage averaged across biological replicate (N=2 for each (DMSO and 10 μM **NV1** treated sample) +/- NAI) at TERC locus. Read coverage is plotted 5' --> 3' (TERC lncRNA). (C) Mutation frequency for DMSO and **NV1** treated samples, comparing mutation rate of NAI (acylating reagent) treated samples relative to DMSO.

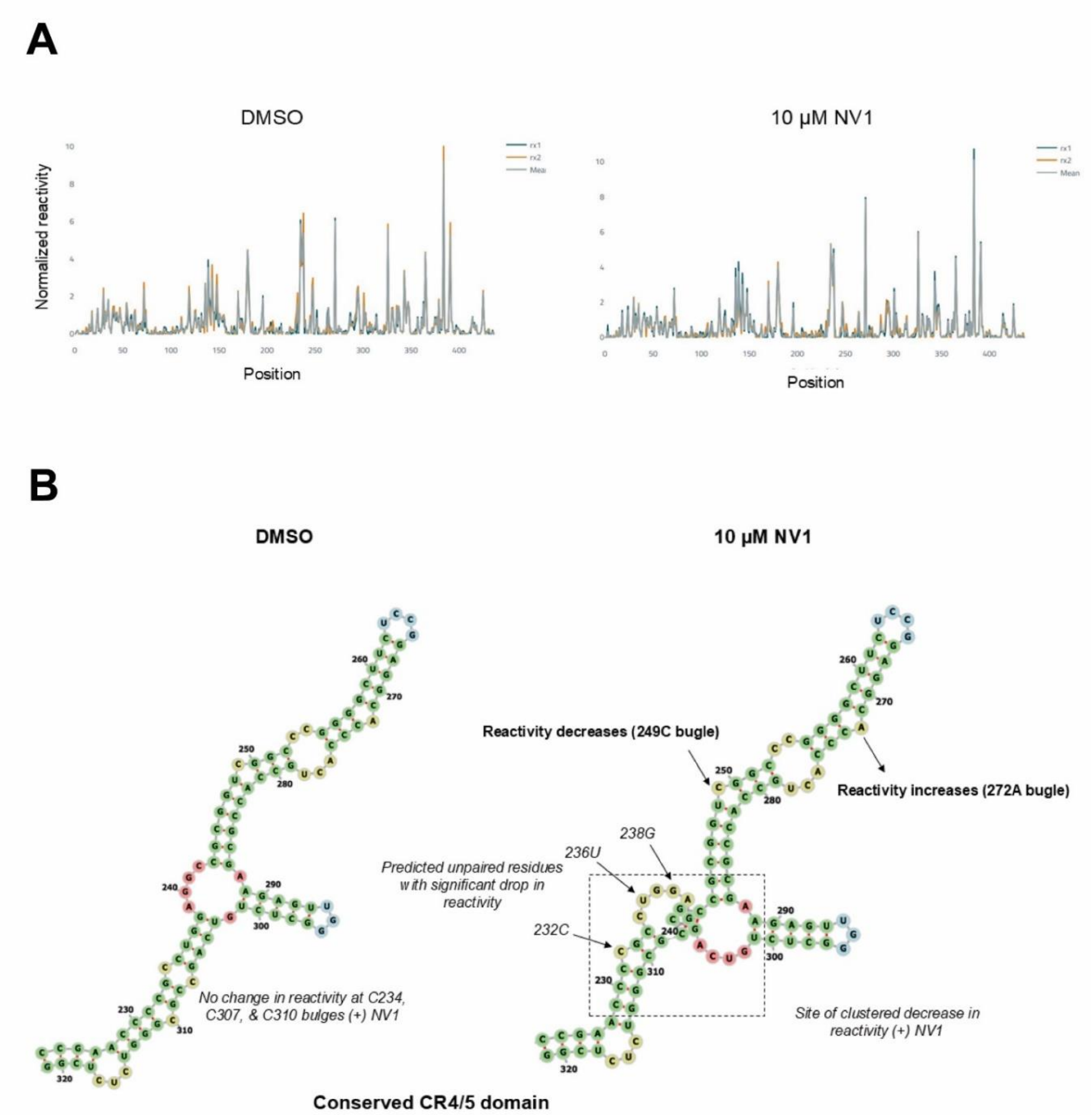


**Supplemental Figure 6. Cellular SHAPE-MaP reactivity profiles and secondary structural renderings of TERC CR4/5 domains with SHAPE constraints for DMSO and NV1 treated samples.** (A) Normalized reactivity profiles (N=2) for DMSO and 10 µM **NV1** treated samples. For each condition, the reactivity trace of each individual replicate is shown with the mean signal (gray) across the two replicates. (B) SHAPE-MaP constrain secondary structure rendering of CR4/5 domain with DMSO (left) or 10 uM NV1 (right) treatment. Boxed region indicates site of significant clustered changes in NAI reactivity in the presence of NV1. The bulged residues 249C is suspected to be important for compound engagement as this residue has the most significant reduction in reactivity. Secondary structures with SHAPE reactivity constraints were visualized using Forna [2]. Nucleotides are colored based on structure (stems = green, multiloops/junctions = red, interior loops = yellow, hairpin loops = blue, 5' & 3' unpaired region = orange).

**a**

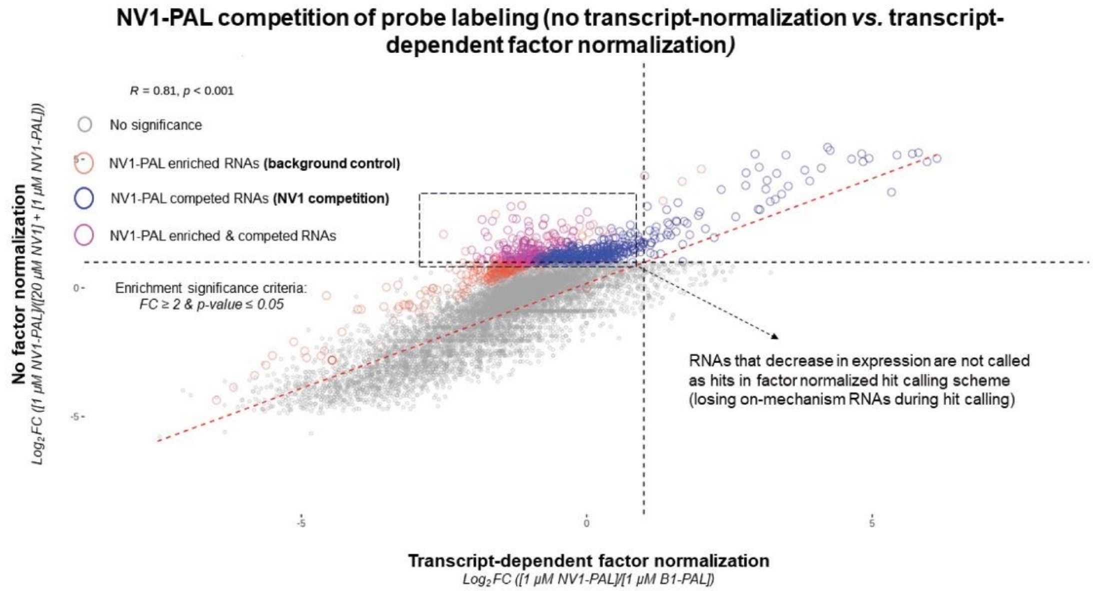


**b**

**Sample code for Implementation of RNA-dependent factor normalization using DESeq2 to normalize read counts of probe-enriched RNAs by covariation in transcript abundance across samples:**

**1)** Create read count table from RNAseq .bedfiles to scale covariation in read counts of targeted enrichment (chem-CLIP) studies

**2)** Define normalization matrix in DESeq object for factor normalization

**3)** Set size factors or assign to default & run differential enrichment analysis

```
for (k in 1:dim(normalization_matrix)[2]) {
            normalization_matrix1[,k] <- normalization_matrix[,k] + 1
}           #add one to each read count value per transcript
normalization_matrix2 <- normalization_matrix1 / exp(rowMeans(log(normalization_matrix1)))
            #divide out geometric mean for each transcript
group <- factor(rep(1:3,each=2))
condition <- factor(rep(rep(c("sample","control"),each=3)))
d <- data.frame(group, condition)
as.data.frame(d)
dds <- DESeqDataSetFromMatrix(countData=round(chem-CLIP_matrix),
                    colData=d,
                    design=~condition,
)
            #define chem-CLIP matrix with raw read counts per transcript
normalizationFactors(dds) <- as.matrix(normalization_matrix2)
            #define normalization matrix in DESeq2 object for scaling differential enrichment
dds <- DESeq(dds)
            #construct object with normalized read count values that can be used as input for differential enrichment analysis
```

**c**

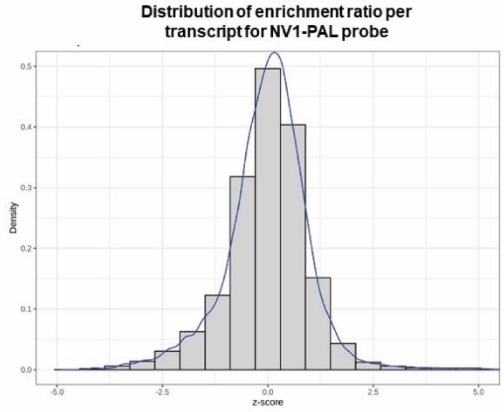


**Z-score calculation:**

1) Enrichment ratio$_{transcript}$ = Log2(chem-CLIP (TPM+1) $_{transcript}$ / RNAseq (TPM+1) $_{transcript}$)

2) Filter for probe-enrichment (TPM) > 0

3) Calculate z-score based on filtered RNAs (~17k RNAs) and assign cut-off for hit calling strategy

**Supplemental Figure 7. Alternative hit calling strategies explored using transcript-dependent factor normalization & z-score analysis.** (A) Scatterplot of $\log_2$ fold-change of 1 μM **NV1-PAL** without or with 20 μM **NV1** without (y-axis) or with (x-axis) factor normalization**.** In this strategy, RNAs with a higher abundance relative to the control sample are not called as hits regardless of degree of probe enrichment/labeling competition. (B) Example of R code to factor scale probe enrichment/degree of competition of probe labeling using parallel RNAseq analysis. Raw read counts from the bulk RNAseq are transformed and defined as the normalization matrix in the DESeq2 object prior to running the differential enrichment analysis. (C) An enrichment ratio is calculated ($\log_2$ (chem-CLIP$_{TPM+1}$ /RNAseq$_{TPM+1}$)) with the assumption that specific or high-affinity compound RNA targets will have an increased TPM relative to bulk RNAseq. Z-score thresholds can be set and compared to DESeq2 output. These strategies did not improve hit calling efficiency.

**A**

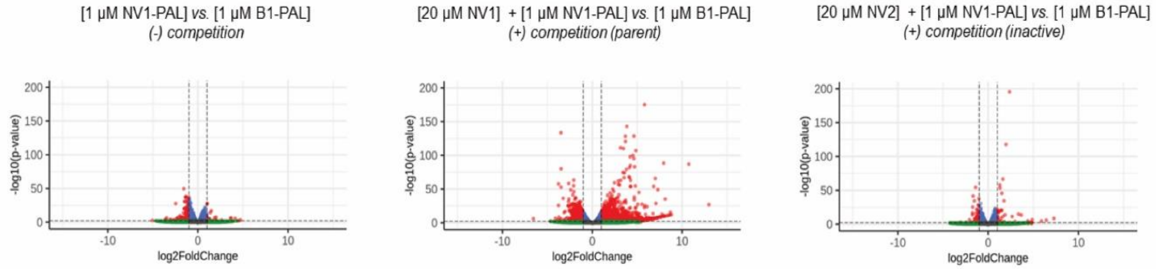


| Expression change | NV1-PAL (-) competition | NV1-PAL (+) competition (parent) | NV1-PAL (+) competition (inactive) |
|---|---|---|---|
| UP | 108 | 2197 | 105 |
| DOWN | 338 | 1387 | 156 |

**9161 total differentially spliced genes in NV-8 PAL treated samples (+/- competition)**

**B**

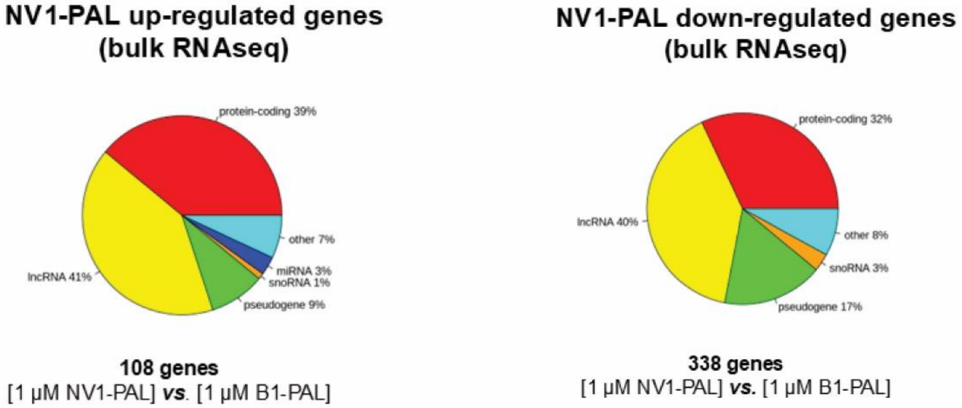


**Supplemental Figure 8. Bulk RNAseq expression profiling of parallel processed NV1-PAL treated samples.** (A) Volcano plots of **NV1-PAL** treated samples (left), **NV1-PAL** with 20 µM **NV1** samples (middle), **NV1-PAL** with 20 µM **NV2** samples (right) *vs.* B1-PAL. (B) Distribution of biotypes of up-regulated & down-regulated RNAs following **NV1-PAL** relative to **B1-PAL** treatments.

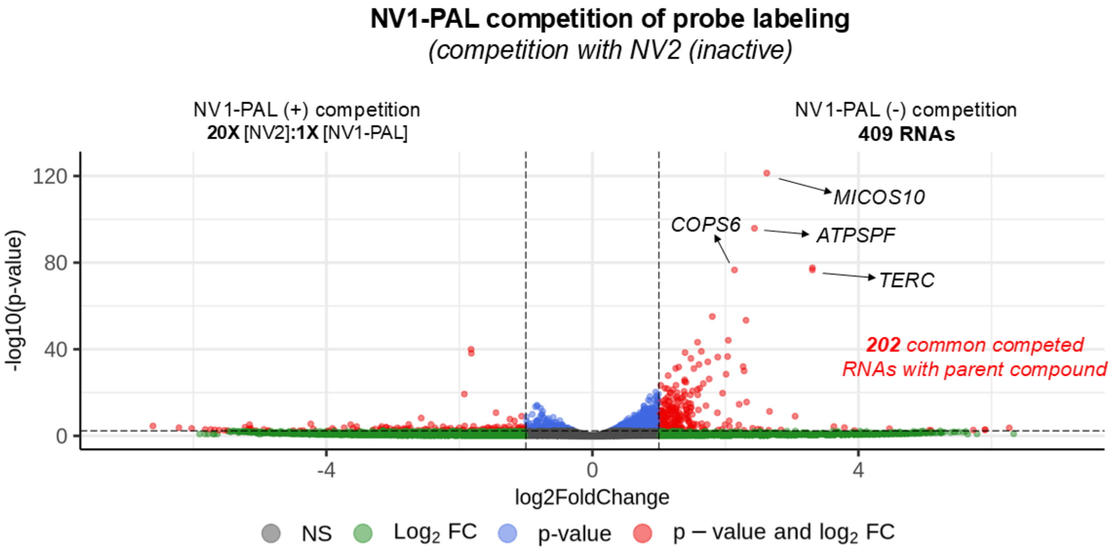


**Supplemental Figure 9. Volcano plot of 1 µM NV1-PAL without or with 20 µM NV2.** Volcano plot of $log_2$FC of **NV1-PAL** with 0 µM **NV2** (right quadrant) *vs.* **NV1-PAL** with 20 µM **NV2** (left quadrant). The RNAs in the right quadrant represent higher labeling in the **NV1-PAL** with 0 µM **NV2** samples, indicating loss of probe labeling in the presence of **NV2**. The strongest RNAs competed by **NV1** and enriched over **B1-PAL** were also present (COPS6, MICOS10, TERC) in the **NV2** comparison.

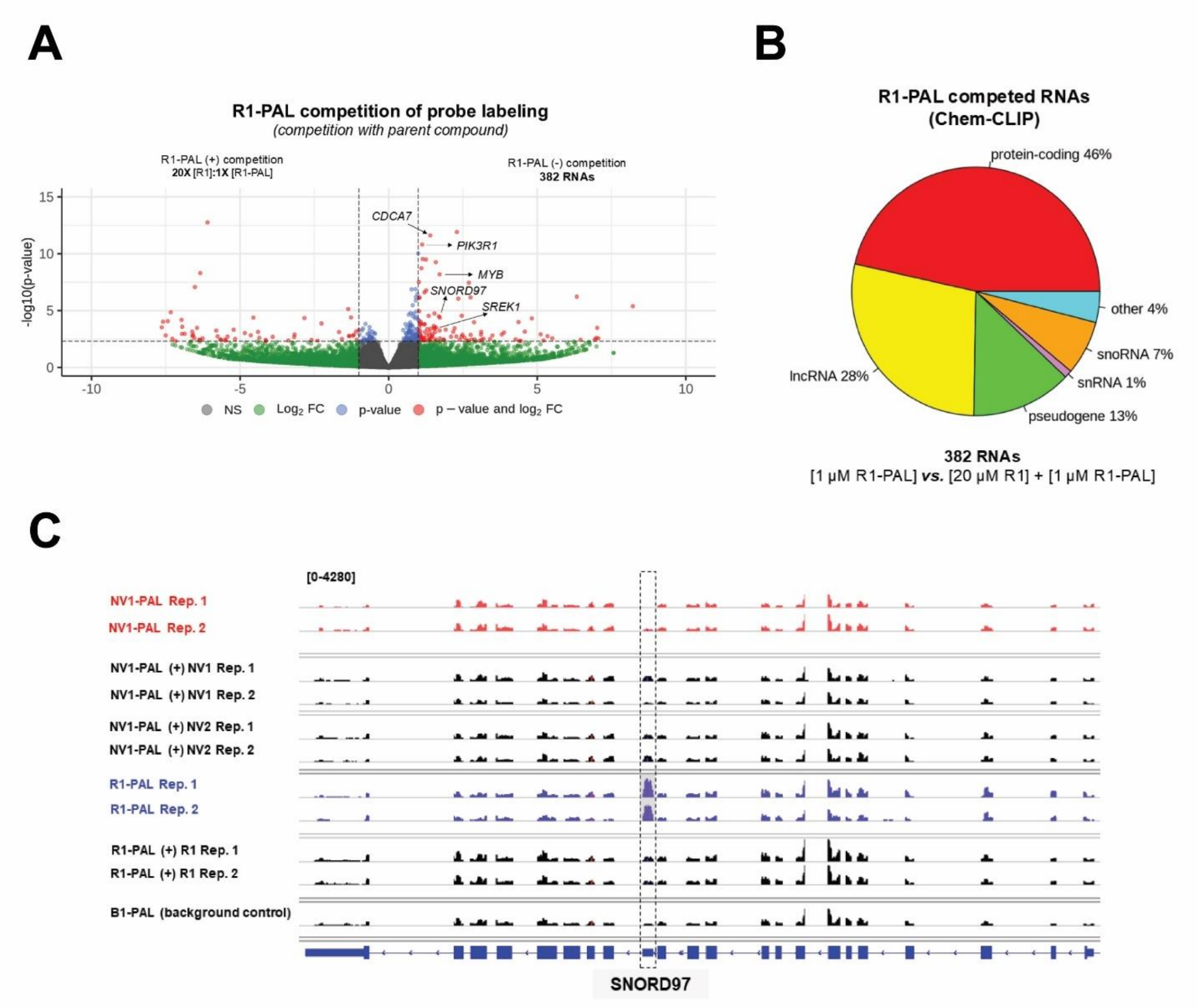


**D**

| Transcript | ΔPSI | 3' nGA exon ending | ΔExpression | Coordinates of cryptic exon |
|---|---|---|---|---|
| CDCA7 | -16.5709 | AGA | Down | chr2:173367832-173368070:+ |
| SREK1 | 3.27331 | AGA | None | chr5:66164608-66164677:+ |
| PIK3R1 | 3.545926 | AGA | Down | chr5:68242956-68243145:+ |
| INTS9 | 3.522479 | AGA | None | chr8:28776167-28776203:- |
| BTBD10 | 3.107866 | AGA | Down | chr11:13419277-13419343:- |
| PITPNB | 1.482293 | UGA | Down | chr22:27894376-27894422:- |
| PDS5B | 2.031735 | AGA | Down | chr13:32688881-32689021:+ |
| MYB | 3.928389 | UGA | Down | chr6:135199526-135199581:+ |
| NSUN4 | 3.164536 | GGA | None | chr1:46357576-46357659:+ |

**Supplemental Figure 10. Differential enrichment of Risdiplam-PAL.** (A) Volcano plot of $log_2FC$ of R1-PAL (-) competition (right quadrant) *vs.* R1-PAL (+) R1 (active) competition (left quadrant). The

RNAs in the right quadrant represent higher labeling in the R1-PAL (-) competition samples, indicating loss of probe labeling in the presence of R1 competitor. The strongest RNAs competed by R1 include MYB, CDCA7, PIK3R1, SREK1, and SNORD97 (snoRNA). (B) Biotype distribution of R1 competed RNAs. (C) Read pile-up at SNORD97 RNA (strongly enriched R1-PAL hit. The exon corresponding to SNORD97 is demarcated by the dotted boxed region. Increased read pile-up in the R1-PAL (-) competition samples relative to the other PAL conditions. Enrichment is depleted in the R1-PAL (+) R1competition sample. (D) Table of 9 RNAs that are R1 competed and differentially spliced (+) R1 with 3' nGA ending exon. Change in percent spliced in (ΔPSI) is relative to differential splicing analysis with B1-PAL ($\log_2$([1 μM R1-PAL/[1 μM and B1-PAL])). A positive PSI indicates increase in splicing inclusion, while a negative value indicates a reduction in splicing. Δ transcript abundance is R1 (+) competition ([20 μM R1] + [1 μM R1-PAL]) *vs.* [1 μM B1-PAL]. The coordinates for the site of the cryptic 3' nGA ending exon are provided in the last column of the table (chr, start, end, strand orientation (+) or (-)).

**Section 2: Synthesis & characterization of chemical photo-probes**

**Compound NV1-PAL (branaplam series-based photo-probe):**

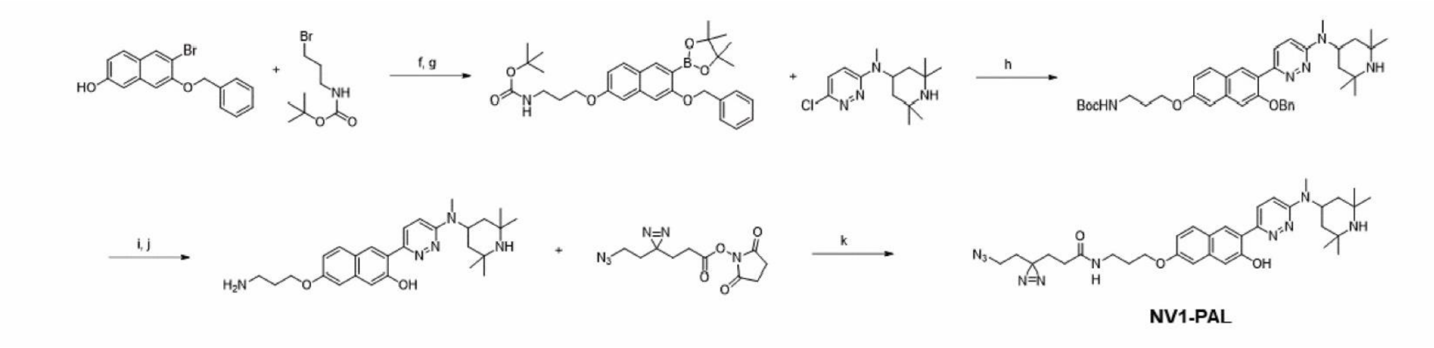


**NV1-PAL:** 7-(3-aminopropoxy)-3-(6-(methyl(2,2,6,6-tetramethylpiperidin-4-yl)amino)pyridazin-3-yl)naphthalen-2-ol (30 mg, 0.056 mmol) and 2,5-dioxopyrrolidin-1-yl 3-(3-(2-azidoethyl)-3H-diazirin-3-yl)propanoate (14.1 mg, 0.05 mmol) were dissolved in DMF (2mL), DIPEA (29 uL, 0.168 mmol) was added to the solution. The mixture was stirred for 1.5 hours at room temperature. LCMS indicated completion of the reaction. The mixture was diluted in MeOH and purified by RP-HPLC using ammonia hydroxide as modifier. desired peak tailed a lot under basic HPLC, since it was clean reaction, desired peaks were not overlapped with impurity. All fractions with desired mass were collected, concentrated to afford 22 mg product as a pale yellow solid. $^1$H NMR (400 MHz, DMSO) δ 13.39 (s, 1H), 8.42 (s, 1H), 8.34 (d, *J* = 9.9 Hz, 1H), 7.98 (t, *J* = 5.6 Hz, 1H), 7.78 (d, *J* = 9.1 Hz, 1H), 7.42 (d, *J* = 9.8 Hz, 1H), 7.18 (s, 1H), 7.10 (d, *J* = 2.4 Hz, 1H), 6.96 (dd, *J* = 8.8, 2.4 Hz, 1H), 5.04 (s, 1H), 4.11 (t, *J* = 6.3 Hz, 2H), 3.23 (q, *J* = 6.8 Hz, 4H), 2.98 (s, 3H), 2.19 (t, *J* = 7.4 Hz, 1H), 1.92 (t, *J* = 7.5 Hz, 4H), 1.72 – 1.60 (m, 6H), 1.24 (m, 16H). LC/MS: retention time: 1.63 min, $[M+1]^+$=629.4, 98% purity; HRMS (m/z): $[M+1]^+$ calcd. for, 629.3676; found, 629.3718;

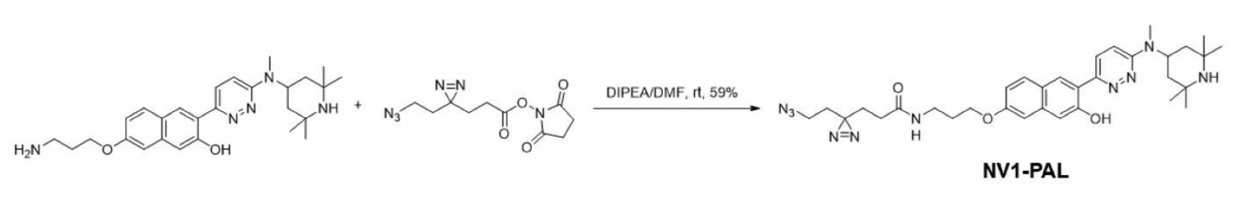

$^{1}H$ NMR (400 MHz, DMSO) δ 13.39 (s, 1H), 8.42 (s, 1H), 8.34 (d, $J$ = 9.9 Hz, 1H), 7.98 (t, $J$ = 5.6 Hz, 1H), 7.78 (d, $J$ = 9.1 Hz, 1H), 7.42 (d, $J$ = 9.8 Hz, 1H), 7.18 (s, 1H), 7.10 (d, $J$ = 2.4 Hz, 1H), 6.96 (dd, $J$ = 8.8, 2.4 Hz, 1H), 5.04 (s, 1H), 4.11 (t, $J$ = 6.3 Hz, 2H), 3.23 (q, $J$ = 6.8 Hz, 4H), 2.98 (s, 3H), 2.19 (t, $J$ = 7.4 Hz, 1H), 1.92 (t, $J$ = 7.5 Hz, 4H), 1.72 – 1.60 (m, 6H), 1.24 (m, 16H).

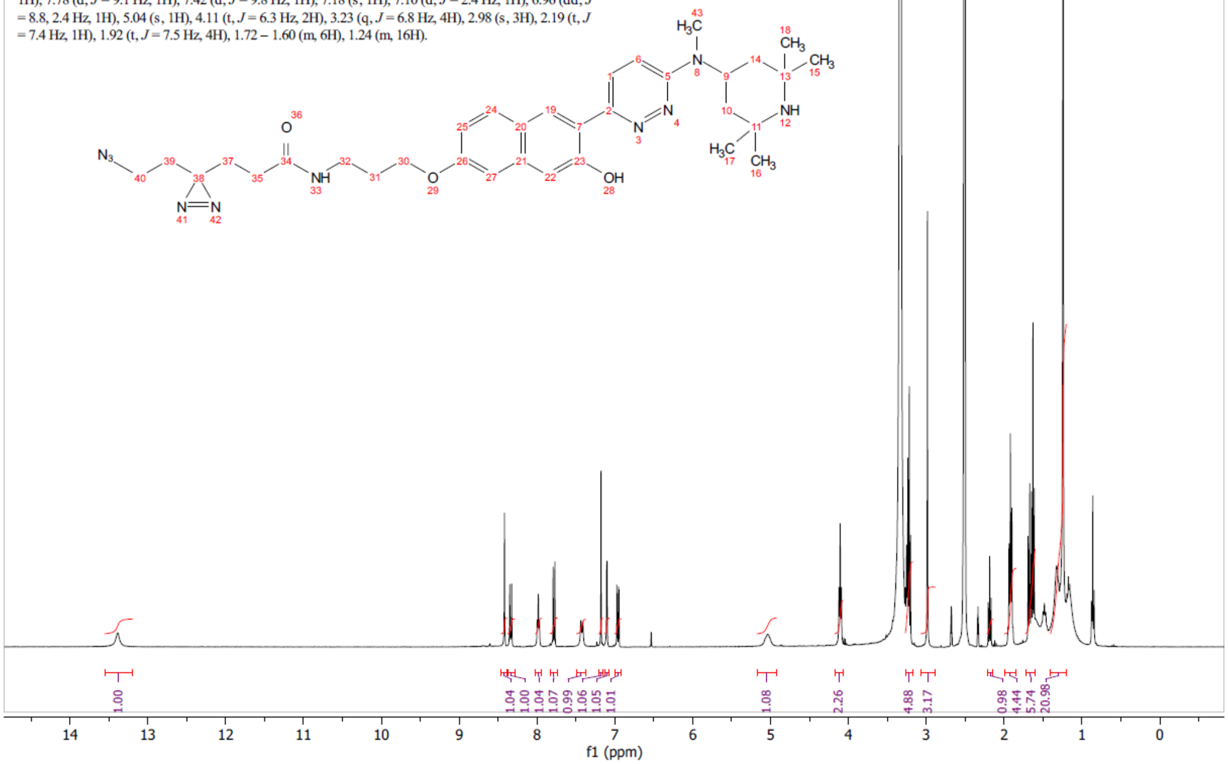


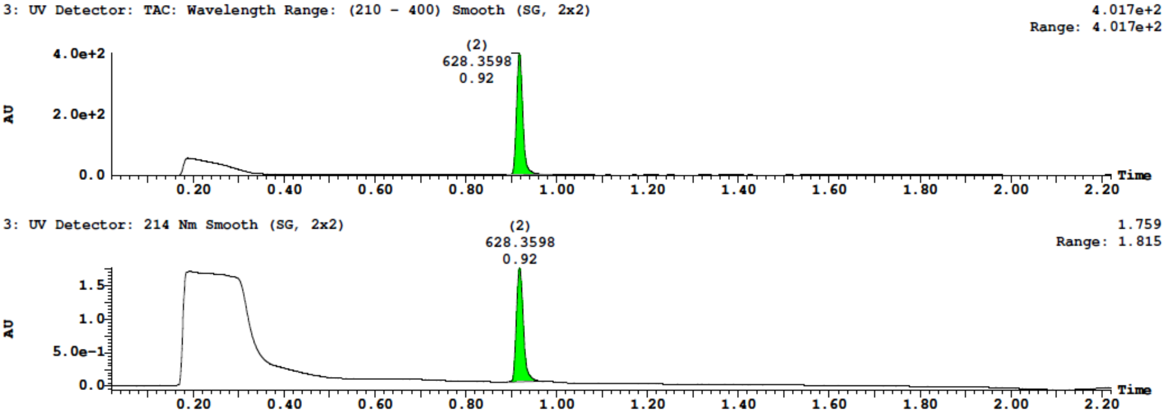


| Peak Number | Time | Width | AreaAbs | Area % | BPM | Mass Found | Conc. |
|---|---|---|---|---|---|---|---|
| 2 | 0.92 | 0.073 | 30256 | 100.00 | 629.4, 281.1 | 628.36 | --- |

| Peak ID | Time | Mass Found | BPM | State | Compound |
|---|---|---|---|---|---|
| 2 | 0.91 | 1257.73,629.37 | 629.4 | OK | Found |

2: (Time: 0.91)

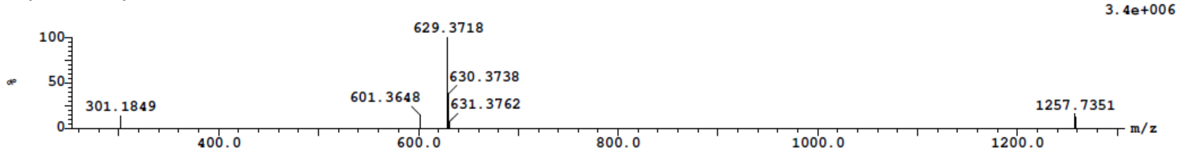

**Compound R1-PAL (risdiplam-based photo-probe):**

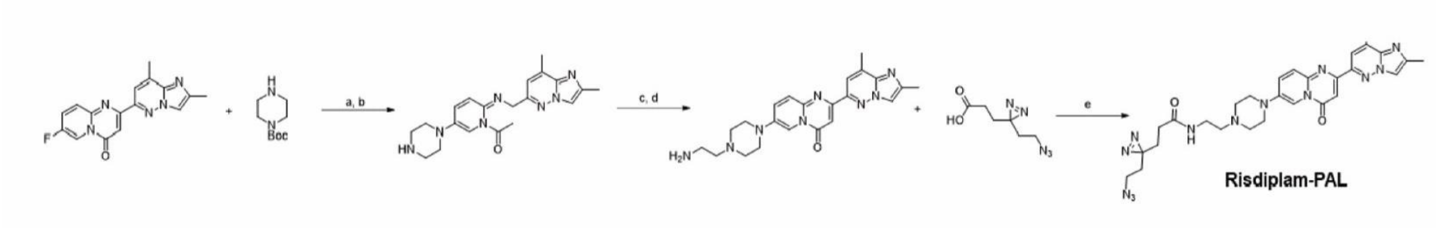


**Risdiplam-PAL**: 3-(3-(2-azidoethyl)-3H-diazirin-3-yl)propanoic acid (15.6 mg, 0.085 mmol) and 7-(4-(2-aminoethyl)piperazin-1-yl)-2-(2,8-dimethylimidazo[1,2-b]pyridazin-6-yl)-4H-pyrido[1,2-a]pyrimidin-4-one (35 mg, 0.071 mmol) were dissolved in DMF (4 mL), DIPEA (50 uL, 0.284 mmol) was added to the solution followed by HATU (54 mg, 0.142 mmol), the mixture was stirred at room temperature for 2 hours, and LC/MS was performed to confirm completion of the reaction. MeOH was added to the reaction mixture, the crude was purified by RP-HPLC, and clean fractions were combined to afford 20 mg yellow solid as product in 40% yield. $^{1}$H NMR conditions: (400 MHz, DMSO) δ 8.31 (d, *J* = 2.7 Hz, 1H), 8.22 – 8.11 (m, 2H), 7.97 (s, 1H), 7.88 – 7.78 (m, 2H), 7.09 (s, 1H), 3.28 (d, *J* = 10.5 Hz, 4H), 3.22 (q, *J* = 6.6 Hz, 4H), 2.64 (s, 3H), 2.60 (t, *J* = 4.9 Hz, 4H), 2.43 (s, 5H), 1.92 (dd, *J* = 8.5, 6.7 Hz, 2H), 1.72 – 1.55 (m, 4H); LC/MS: retention time: 1.05 min, $[M+1]^+$=584.2, 99% purity; HRMS (m/z): $[M+1]^+$ calcd. for, 584.2953; found, 584.2989;

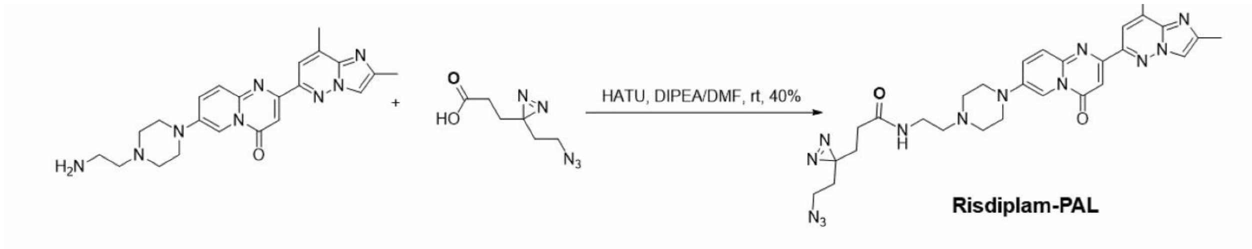

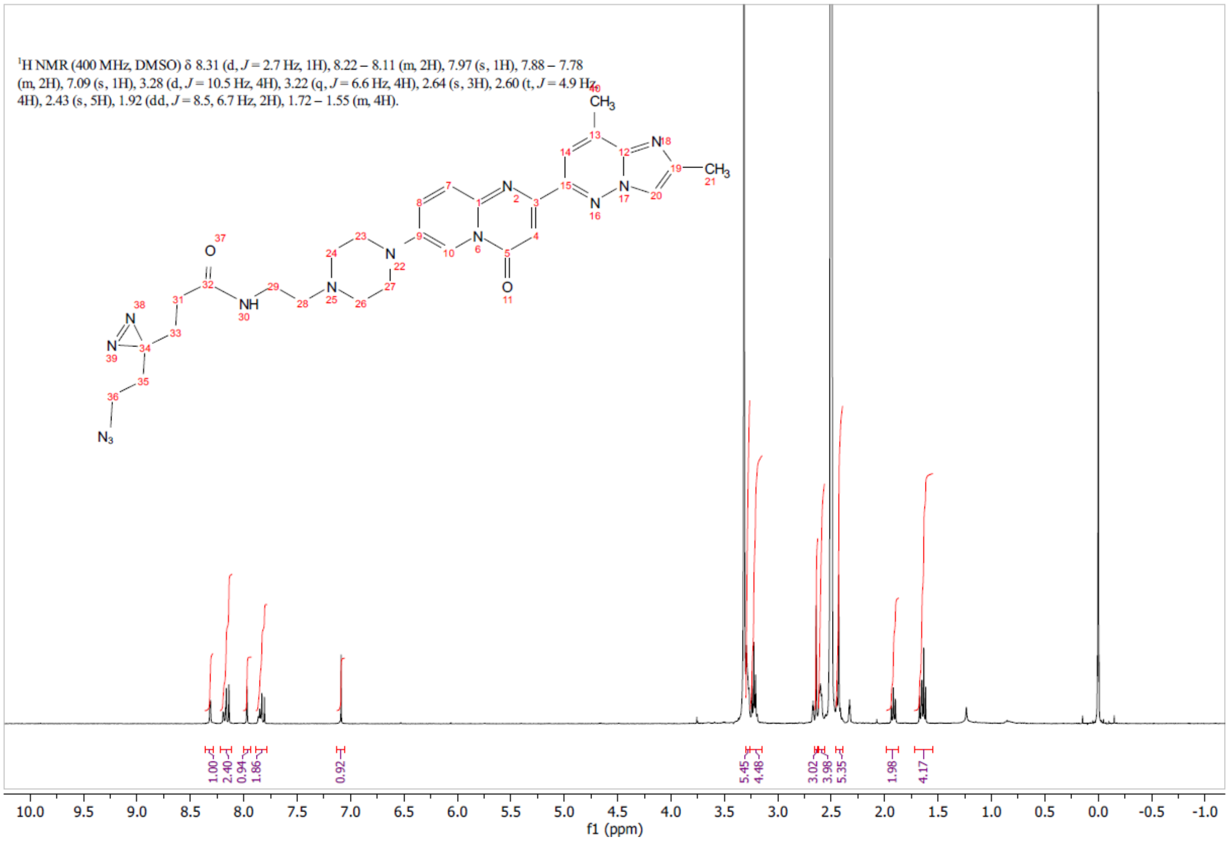


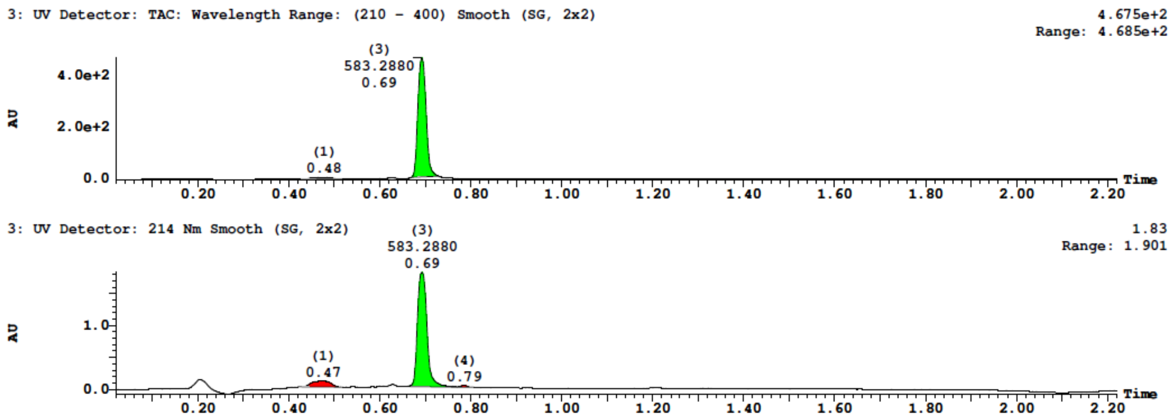


| Peak Number | Time | Width | AreaAbs | Area % | BPM | Mass Found | Conc. |
|---|---|---|---|---|---|---|---|
| 1 | 0.47 | 0.098 | 4582 | 9.42 | 485.4, 211.1 | Not Found | --- |
| 3 | 0.69 | 0.124 | 43493 | 89.43 | 584.3, 315.1 | 583.29 | --- |
| 4 | 0.79 | 0.038 | 558 | 1.15 | 411.2, 211.1 | Not Found | --- |

| Peak ID | Time | Mass Found | BPM | State | Compound |
|---|---|---|---|---|---|
| 3 | 0.69 | 292.65,584.30 | 584.3 | OK | Found |

3: (Time: 0.69)

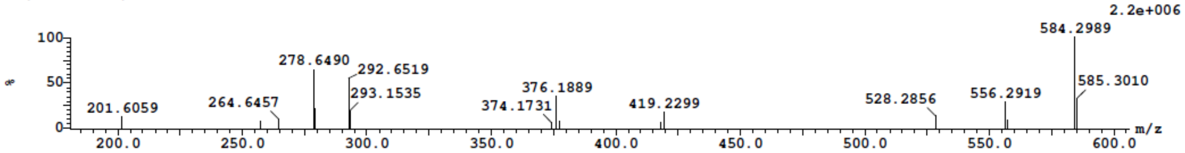


| Mass | Calc. Mass | mDa | PPM | DBE | Formula | i-FIT (norm) |
|---|---|---|---|---|---|---|
| 584.2989 | 584.3017 | -2.8 | -4.8 | 9.5 | C21 H38 N13 O7 | 0.1 |
| 584.2989 | 584.2990 | -0.1 | -0.2 | 10.5 | C17 H34 N19 O5 | 2.6 |
| 584.2989 | 584.3004 | -1.5 | -2.6 | 4.5 | C20 H42 N9 O11 | 4.2 |
| 584.2989 | 584.2977 | 1.2 | 2.1 | 5.5 | C16 H38 N15 O9 | 5.6 |
| 584.2989 | 584.2985 | 0.4 | 0.7 | 17.5 | C32 H38 N7 O4 | 6.0 |
| 584.2989 | 584.2972 | 1.7 | 2.9 | 12.5 | C31 H42 N3 O8 | 6.2 |
